# A Tissue Culture–Free Genome Editing Strategy in Plants Using Broad-Host-Range Viral Vectors Derived from Geminiviruses

**DOI:** 10.64898/2026.02.15.705632

**Authors:** Jitendra Kumar, Anshu Alok, Jack Fox, Ashish Srivastava, Daniel F. Voytas, Feng Zhang, Shahryar F. Kianian

## Abstract

The use of viral vectors offers a promising alternative to traditional transformation methods for creating gene-edited plants. In this study, we developed a novel plant genome editing system by delivering Cas9, Cas12f, and Cas12j nucleases along with their guide RNAs using a broad-host-range geminivirus, *Wheat dwarf India virus* (WDIV), in combination with Ageratum yellow leaf curl betasatellite (AYLCB). Cas9, Cas12f, and Cas12j nucleases were efficiently expressed along with corresponding guide RNAs under viral promoters. By leveraging tRNA spacers in place of external promoters and terminators, we significantly reduced the overall cargo size, streamlining vector design. Additionally, we compared the traditional AtU6-driven gRNA delivery with a novel spacer:gRNA:spacer format in Cas9-expressing lines and observed comparable editing efficiencies. The broad host range of WDIV and AYLCB, combined with this tissue culture–free genome editing platform, opens up possibilities for editing across a wide range of plant species.

## Introduction

Efficient delivery of CRISPR–Cas reagents remains a significant bottleneck in the application of genome editing for crop improvement. Current methods are largely restricted to a limited number of crops and genotypes that are amenable to transformation and regeneration. Viral vectors offer a promising alternative, providing enhanced editing efficiency and the ability to target multiple genes simultaneously (Mikhaylova, 2025; Steinberger and Voytas, 2025). Notably, viral systems hold potential for tissue culture–free and heritable genome editing (Mikhaylova, 2025; Steinberger and Voytas, 2025; Zhang et al., 2022). However, their utility has been limited by a constrained cargo capacity and a narrow host range (Mikhaylova, 2025; Shen et al., 2024). Most viral vector–based editing studies have relied on Cas transgenic plants due to limited cargo capacity, which restricts the method to species and genotypes that can be genetically transformed (Mikhaylova, 2025; Nasti and Voytas, 2021; Zhang et al., 2022).

The discovery of miniature type V CRISPR nucleases such as Cas12f (also known as Cas mini) and Cas12j (CasΦ), which are less than half the size of Cas9, opens new possibilities for viral delivery of both Cas proteins and guide RNAs (Pausch et al., 2020; Xu et al., 2021; Wu et al., 2021). Recent studies have demonstrated the genome editing capabilities of Cas12f and Cas12j in both plant and animal systems (Bigelyte et al., 2021; Kim et al., 2022; Pausch et al., 2020; Wu et al., 2021; Xu et al., 2021; Sukegawa et al., 2023). These enzymes recognize T-rich PAM sequences, characteristic of type V nucleases, and mediate DNA cleavage through the formation of asymmetric homodimers (Takeda et al., 2021; Xiao et al., 2021; Pausch et al., 2021).

Geminivirus-based vectors provide a robust platform for the delivery of genome editing reagents and donor DNA (Baltes et al., 2014; Catoni et al., 2018; Gong et al., 2024). Deconstructed geminiviral replicons have been widely used for the transient expression of CRISPR components. However, these DNA replicon systems are limited by poor systemic movement, restricting expression to inoculated tissues (Baltes et al., 2014; Acha et al., 2021; Gong et al., 2024). To date, no geminivirus-based system has been reported that enables the delivery of both the Cas nuclease and guide RNA while maintaining systemic spread.

In a previous study, we engineered *Wheat dwarf India virus* (WDIV; family *Geminiviridae*) as a vector for virus-induced gene silencing (VIGS) and genome editing (VIGE). We demonstrated efficient silencing of the PDS gene and CRISPR guide RNA delivery in Cas9-expressing *Nicotiana benthamiana* lines, resulting in targeted insertions and deletions (Kumar et al., 2024). WDIV is a unique leafhopper-transmitted geminivirus with a broad host range that includes both monocots and dicots such as wheat, oat, barley, maize, soybean, tobacco, and sugarcane. It is also associated with both alpha-and beta-satellites (Kumar et al., 2012, 2014, 2020a, 2024). Both of these features make WDIV a unique virus and suitable for the VIGE vector.

In this study, we leveraged the smaller genome size of WDIV-associated Ageratum yellow leaf curl betasatellite (AYLCB). We engineered it for the delivery of both CRISPR guide RNAs and Cas effectors in plant tissues. As a proof of concept, we first validated the vector’s systemic delivery and expression potential using an AmCyan reporter. Following successful expression of AmCyan, we optimized the vector to reduce the overall size of CRISPR constructs by replacing external promoters and terminators with transfer RNA (tRNA) spacers and expressing gRNA under the control of the viral promoter. We observed similar editing efficiencies with AtU6-driven gRNA and viral promoter-driven gRNA in Cas9-expressing lines. We then co-infiltrated wild-type *N. benthamiana* leaves with WDIV and AYLCB constructs carrying gRNAs and one of three Cas effectors (Cas9, Cas12f, or Cas12j). Here, we demonstrated a novel approach for a tissue culture–free genome editing platform for high-throughput editing.

## Results

### Features of WDIV and AYLCB promoters

Based on previous studies (Zhang et al., 2012; Shukla et al., 2013), we selected a 455-nucleotide region upstream of the C1 gene on the complementary strand of AYLCB (GenBank accession MN240347) as a candidate promoter (Figure S1). Analysis using the PlantCARE database (Lescot et al., 2002) revealed typical cis-regulatory elements, including TATA-box, CAAT-box, G-box, AE-box, TCA-element, and TC-rich repeats, consistent with earlier reports for other betasatellites.

We also selected the long intergenic region (LIR) of WDIV (GenBank accession MN240329), known for its bidirectional promoter activity in mastreviruses (Boulton, 2002). PlantCARE analysis identified multiple regulatory elements, including the TATA-box, CAAT-box, GATA-motif, AAAG-motif, and G-box (Figure S1), in agreement with previous findings (Xie et al., 2003; Khan et al., 2015; Alok et al., 2019). The presence of these elements strengthens the candidacy of these sequences as promoters to regulate transcription.

### The WDIV complementary strand promoter exhibits strong activity

It was crucial to test the strength of the internal promoters present on the viral vector to use them for driving transcription and minimizing the cargo size. To evaluate the strength of the promoter, transgenic tobacco plants were generated with GUS driven by AYLCB-C1, WDIV-VF, WDIV-CF, or CaMV35S promoters (Figure 1A). GUS staining showed the strongest expression in plants transformed with WDIV-CF, followed by CaMV35S and WDIV-VF, with AYLCB-C1 showing the weakest activity (Figure 1C–F). These results were corroborated by qRT-PCR, which revealed the highest GUS transcript levels in WDIV-CF lines and the lowest in AYLCB-C1 lines (Figure 1G). Comparable expression of the hygromycin resistance gene (hpt) across all transgenic lines (AYLCB-C1, WDIV-VF, WDIV-CF, and CaMV35S) suggested equal copy number of the integrated T-DNA.

**Figure 1.**
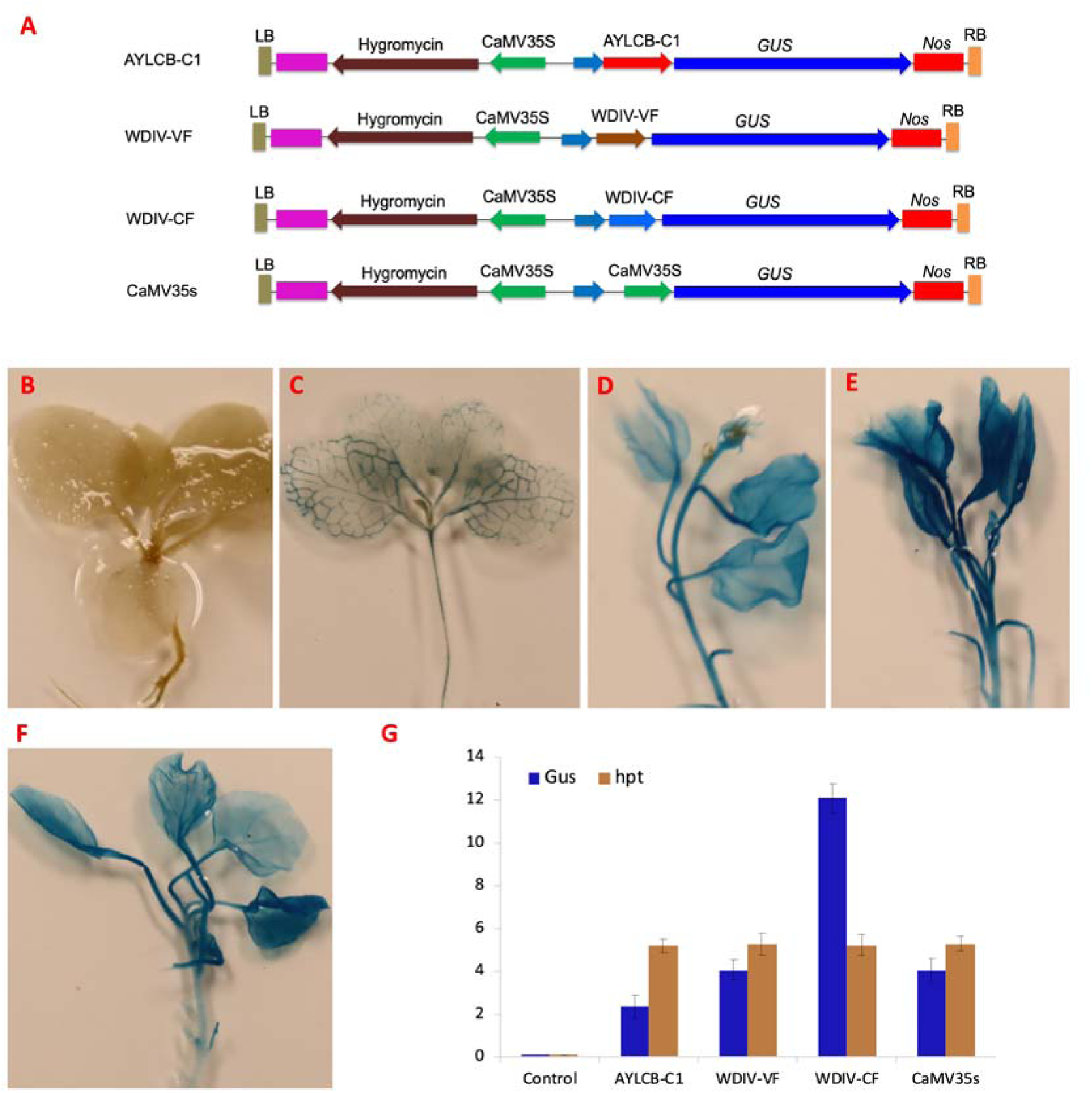
Comparative analysis of promoter activity in transgenic tobacco using GUS expression. (A) Schematic diagrams of pCAMBIA1305 constructs with GUS driven by AYLCB-C1, WDIV-VF, WDIV-CF, or CaMV35S promoters. (B–F) Histochemical GUS staining in leaves and stems of T seedlings: (B) non-transgenic control, (C) AYLCB-C1::GUS, (D) WDIV-VF::GUS, (E) WDIV-CF::GUS, and (F) CaMV35S::GUS. (G) qRT-PCR quantification of GUS and hpt transcript levels. Actin was used as the internal reference. Data represent means ± standard error from three biological replicates.

### AYLCB functions as an efficient viral delivery and expression vector

We next assessed the delivery and expression efficiency of AYLCB-based vectors using *AmCyan* as a visual marker. We tested CaMV35S (35AN), WDIV-VF (VFAN), and WDIV-CF (CFAN) in addition to the AYLCB-C1 (AN) promoter to compare the delivery and expression potential (Figure 2A). Inoculated leaves at 5 days post-inoculation (DPI) showed the strongest *AmCyan* signal in CFAN, followed by 35AN, VFAN, and AN construct (Figure 2B–E) as observed previously with GUS expression. Newly emerged systemic leaves at 21 DPI also showed detectable expression (Figure 2H–K). Control leaves showed weak autofluorescence or no signal (Figure 2F, M).

**Figure 2.**
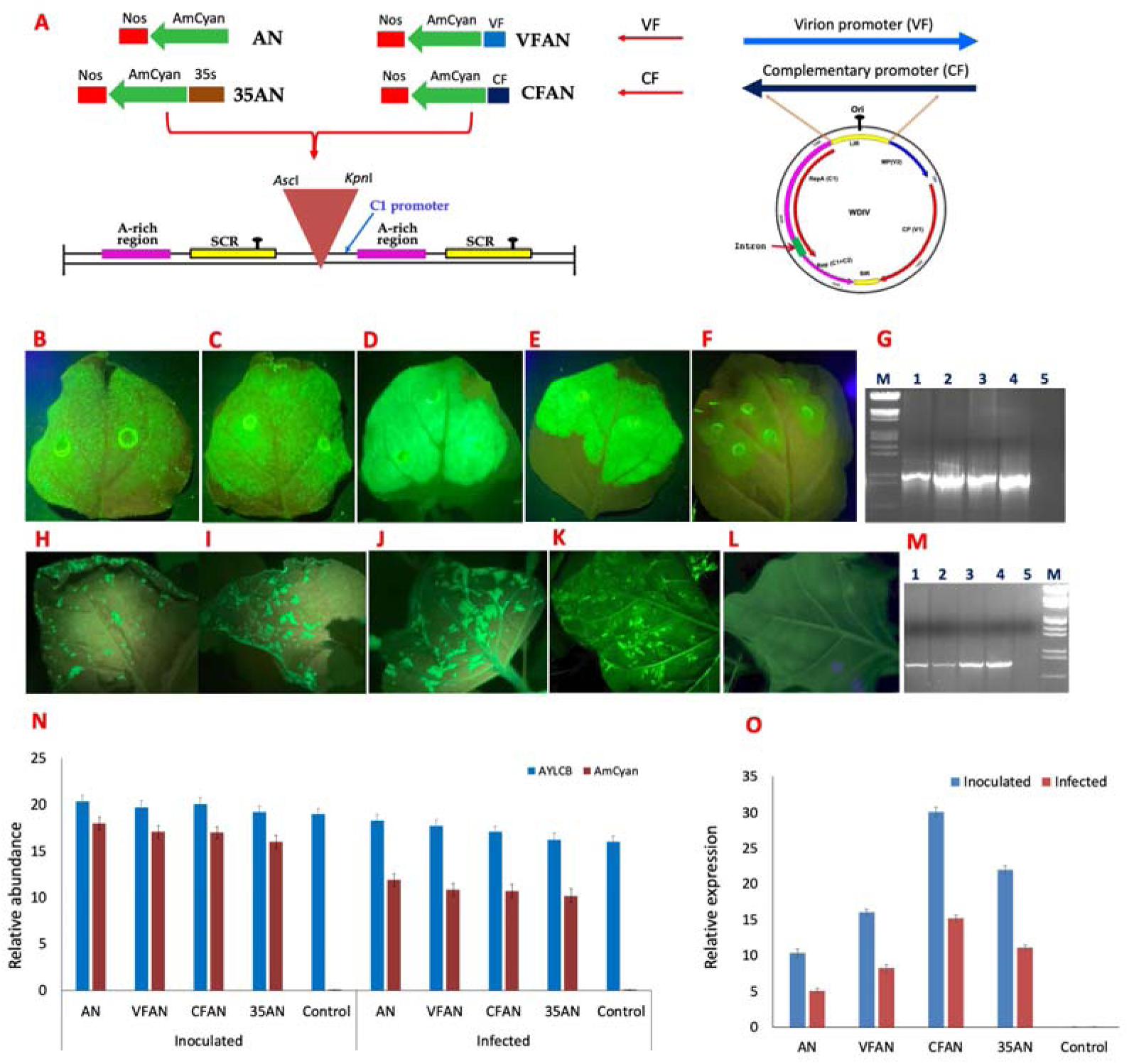
Evaluation of WDIV-derived promoters for systemic expression of AmCyan in tobacco. (A) Schematic of AmCyan constructs with different promoters (AN, 35AN, VFAN, CFAN). (B–F) Fluorescence in inoculated leaves at 5 dpi; (H–L) systemic expression at 3 weeks post-inoculation. (G, M) PCR detection of AmCyan in inoculated and systemic leaves. (N) qPCR analysis of vector and AmCyan DNA accumulation. (O) AmCyan transcript quantification by qRT-PCR. Actin was used as the reference gene. Error bars represent standard error (n = 3).

PCR confirmed *AmCyan* presence in inoculated and systemic tissues (Figure 2G, M). qPCR showed similar levels of AYLCB DNA in both leaf types, while *AmCyan* DNA and transcript levels were reduced in systemic leaves relative to inoculated ones (Figure 2N, O). Across all samples, CFAN drove the highest *AmCyan* expression. This also confirms that the C1 promoter of AYLCB vector can efficiently drive expression of gene cloned on the AYLCB vector.

### Amplicon sequencing confirms genome editing

PCR Amplicons covering target regions were sequenced by Genewiz (Azenta Life Sciences) and analyzed using Cas-Analyzer (Park et al., 2017) and CRISPResso2 (Clement et al., 2019). Both tools produced similar indel frequencies for Cas9-edited samples (Figure S2). For miniature Cas nucleases, CRISPResso2 consistently predicted slightly higher indel frequencies (Figure S3, Table 1, Table S1), which could be due to differences in the parameters used by the two tools. Deletions ranged from 1 to several nucleotides. Cas-Analyzer results are presented in the sections below.

**Table 1:**
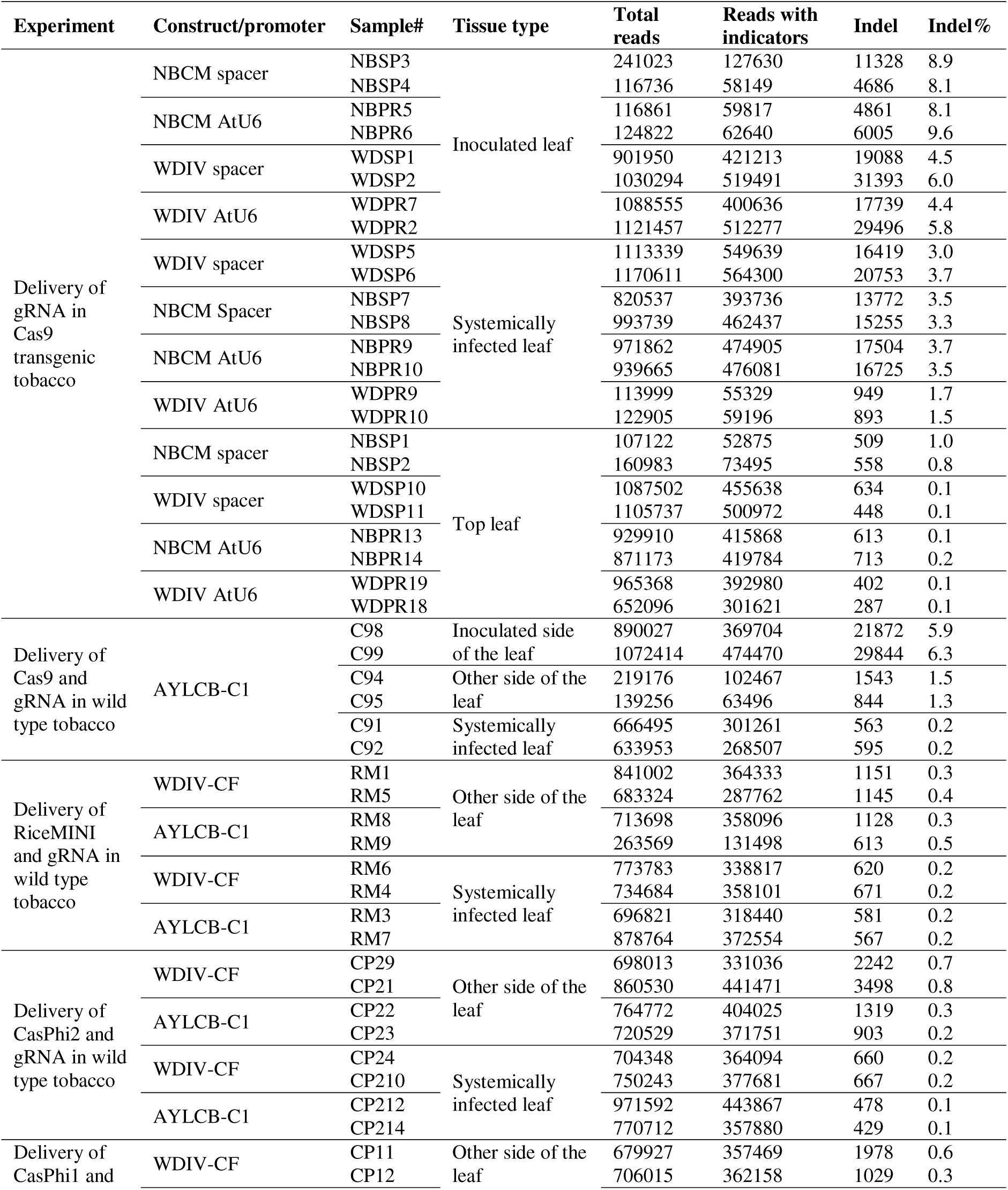
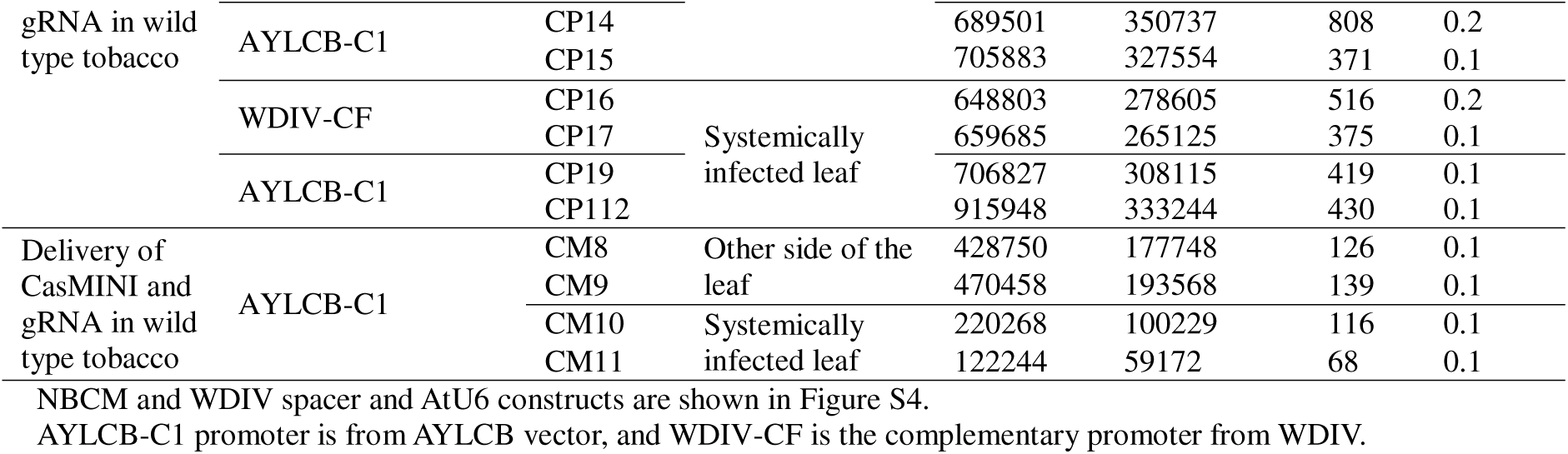
Summary of Indels (insertion and deletion) in different samples.

### Viral promoters support gRNA expression comparable to AtU6

As a first step to drive editing using a viral promoter, we compared gRNA expression driven by viral versus Pol III promoters. We delivered gRNAs using two designs: AtU6::gRNA and spacer::gRNA::spacer, via WDIV and AYLCB vectors (Figure S4). gRNA expression was regulated by the AtU6 promoter in the AtU6::gRNA construct, whereas the viral promoters, AYLCB-C1 or WDIV-VF, regulated the expression in spacer::gRNA::spacer constructs. These constructs are referred to as WDIV AtU6, NBCM AtU6, WDIV spacer, and NBCM spacer (Figure S4, Table 1). Amplicon sequencing showed the highest indel frequencies in inoculated leaves, followed by systemic leaves, with the lowest frequencies in upper leaves (Figure 3, Table 1).

**Figure 3.**
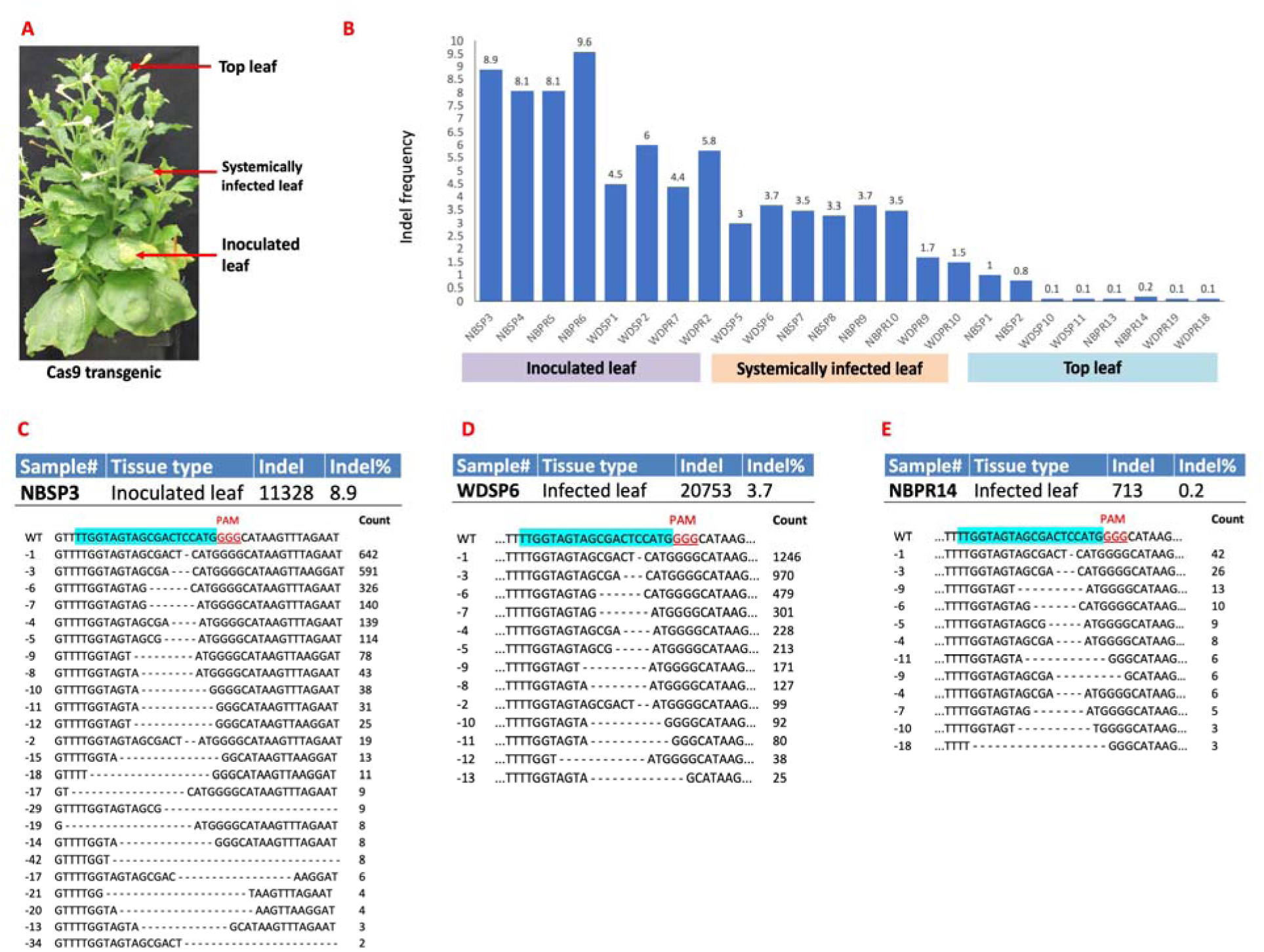
gRNA delivery and genome editing efficiency in Cas9-expressing tobacco. (A) Inoculation and sampling regions in Cas9 transgenic tobacco. (B) Indel frequencies in inoculated, systemic, and upper leaves. (C–E) Representative deletion mutations in the PDS gene. gRNA sequences are shown in turquoise; PAM sites in red. See Tables S2–S4 for additional mutation data.

Importantly, spacer::gRNA::spacer constructs under viral promoters showed editing efficiencies comparable to AtU6::gRNA constructs (Table 1), suggesting that viral promoters can drive functional gRNA expression. Among all treatments, the NBCM spacer produced the highest indel frequencies in top leaves, possibly due to the smaller size (∼250 bp) of the spacer::gRNA::spacer cassette compared to AtU6::gRNA (∼550 bp). Predominant mutations included single-and three-nucleotide deletions (Figure 3C–E, Table S2–S4).

### AYLCB vector enables co-delivery of Cas9 and gRNA

To test Cas9-mediated editing via a single vector, we designed a construct expressing spacer::Cas9::spacer::gRNA::spacer under the AYLCB-C1 promoter (Figure 4A). After co-inoculation with WDIV, leaves were sampled at 12 DPI, and adjacent tissues at 15 DPI (Figure 4B). Amplicon sequencing confirmed editing in all regions, with the highest indel frequencies observed on the inoculated side of the leaf (Figure 4C). Systemic tissues showed reduced editing but still confirmed delivery and expression. Common indel patterns included 1-, 3-, 4-, 6-, 7-, and 9-bp deletions (Figure 4D–F, Table S5). Considering a lower abundance of *AmCyan* DNA and transcripts in the systemically infected leaf tissues (Figure 2N and 2O), a lower editing efficiency in the systemically infected leaves could be attributed to insert instability due to the large size of Cas9.

**Figure 4.**
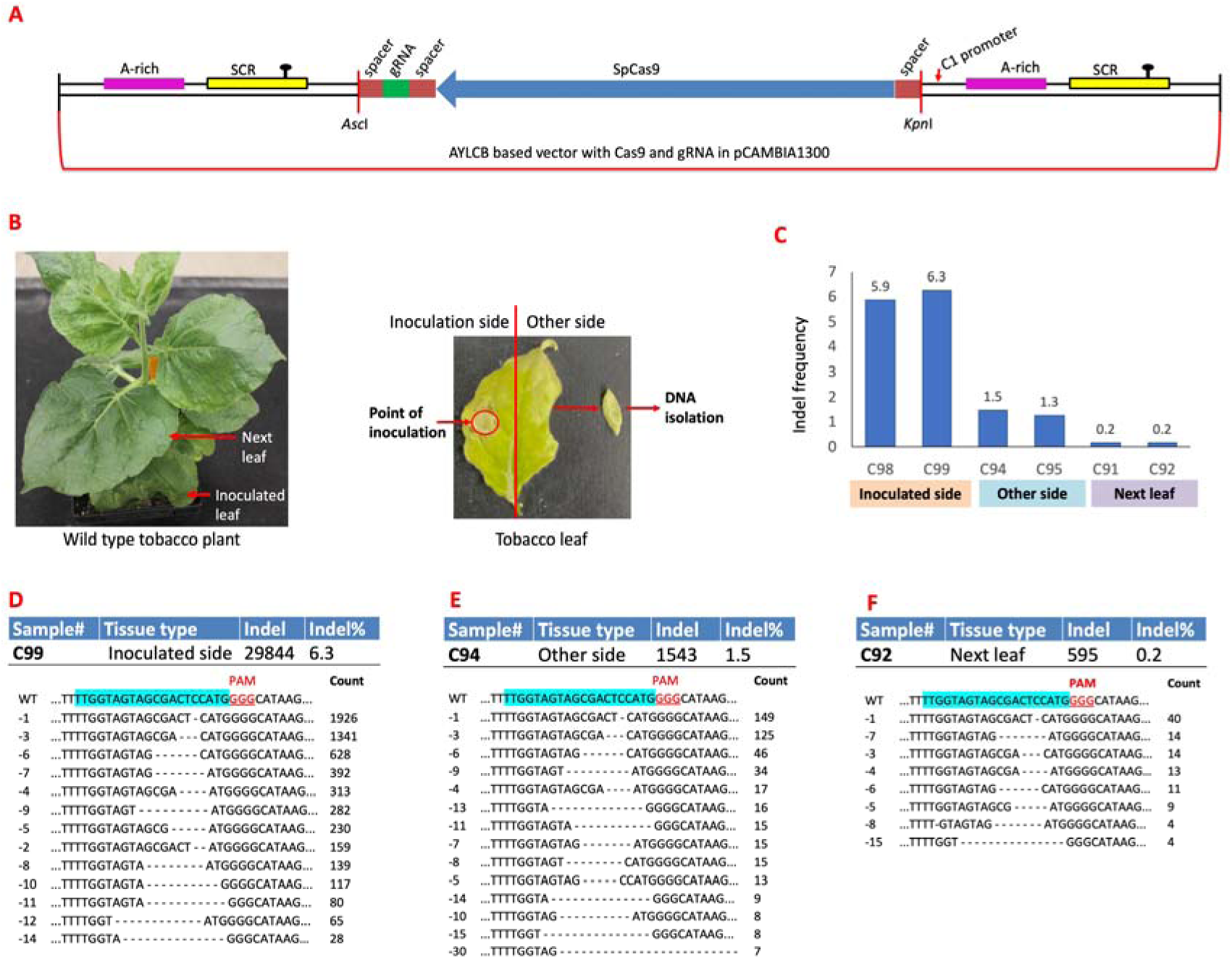
Local and systemic gene editing via delivery of Cas9 and gRNA using the AYLCB vector. (A) Diagram of Cas9/gRNA expression cassette in AYLCB. (B) Sample collection scheme from inoculated and adjacent leaves. (C) Indel frequencies at different tissue locations. (D–F) Examples of deletion mutations in inoculated, opposite, and neighboring leaves. Target and PAM sequences are highlighted. See Table S5 for details.

### AYLCB supports Cas12j (Cas**Φ**1 and Cas**Φ**2) editing

We next tested the delivery of miniature Cas12j effectors, CasΦ1 and CasΦ2, using the spacer::Cas::spacer::gRNA::spacer strategy under the AYLCB-C1 promoter (Figure 5A). Following AYLCB and WDIV co-infection, leaves were sampled and sequenced (Figure 5B).

**Figure 5.**
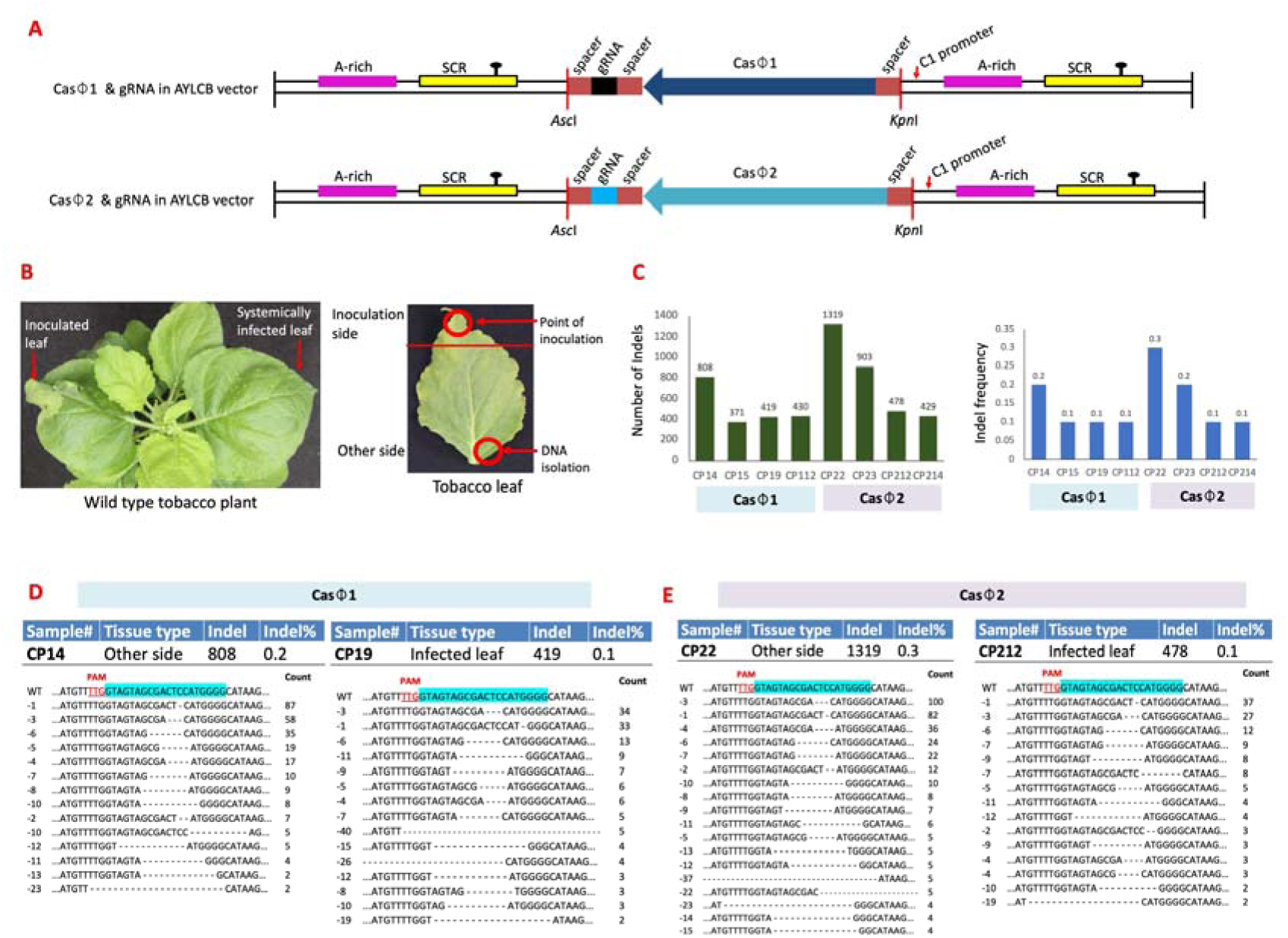
Efficient genome editing by delivery of CasΦ1 and CasΦ2 nucleases with gRNAs. (A) Schematic of CasΦ1 and CasΦ2 constructs in AYLCB. (B) Sample collection strategy from inoculated and systemic leaves. (C) Indel counts and frequencies in distal and systemic tissues (CP series). (D, E) Representative deletions from each tissue type. Detailed mutation data in Tables S6 and S7.

Indel frequencies were higher in tissues on the opposite side of the inoculated leaf compared to systemic leaves (Figure 5C). A range of deletions was detected, whereas both of the Cas effectors had similar editing profiles (Figure 5D, E; Tables S6, S7). Moreover, the Cas12j showed a higher systemic editing efficiency compared to Cas9, which could be attributed to its smaller size and improved delivery.

### Cas12f (CasMINI and RiceMINI) delivery and editing

We then tested Cas12f effectors (CasMINI v3.1 and RiceMINI) using the same strategy (Figure 6A). AYLCB vectors were co-inoculated with WDIV, and tissues were sampled at similar time points (Figure 6B). RiceMINI induced low but detectable indel frequencies, with single-nucleotide deletions being most common in inoculated tissue (Figure 6C–E; Table S8). In contrast, CasMINI yielded minimal editing (Figure 6C; Table S9). A codon-optimized version could be tested to rule out the poor performance of the CasMINI.

**Figure 6.**
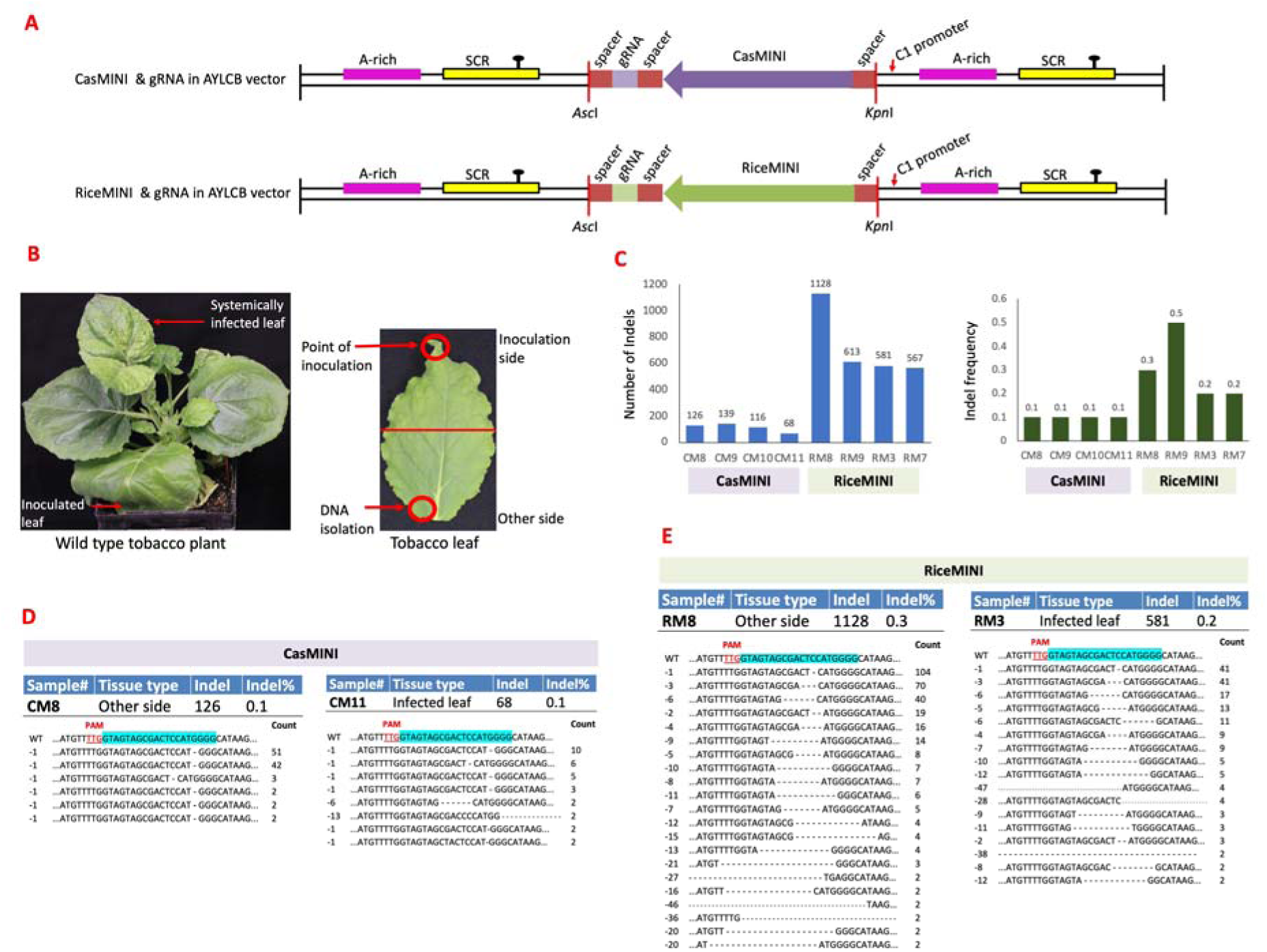
Functional validation of CasMINI and RiceMINI nucleases for genome editing in planta. (A) Construct diagrams of CasMINI and RiceMINI with gRNAs in AYLCB. (B) Leaf samples for editing analysis. (C) Indel frequencies in distal and systemic tissues (CM and RM series). (D, E) Representative mutations. See Tables S8 and S9 for full mutation sets.

### WDIV-CF promoter enhances editing by miniature Cas effectors

Given the strong expression profile of WDIV-CF (Figures 1 and 2), we tested its effect on editing efficiency. The WDIV-CF promoter was inserted upstream of CasΦ1, CasΦ2, and RiceMINI constructs (Figure 7A). Inoculated plants showed increased editing efficiencies, especially in distal tissues (Figure 7B). A broader range of indels was observed, with deletions varying in size (Figure 7C–E; Tables S6–S8). The addition of WDIV-CF significantly enhanced editing compared to constructs using only the AYLCB-C1 promoter (Table 1).

**Figure 7.**
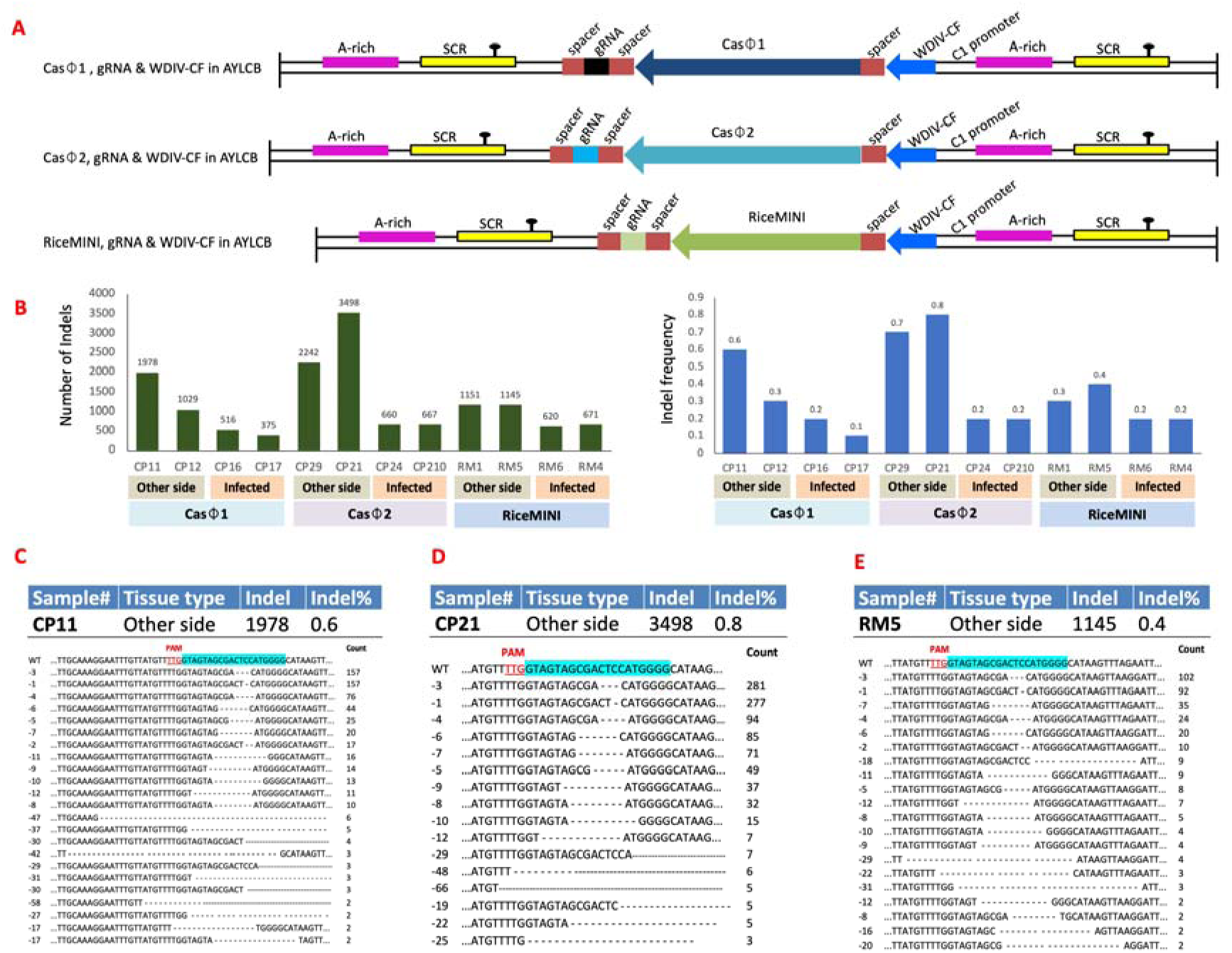
Gene editing using WDIV-CF-driven expression of CasΦ1, CasΦ2, RiceMINI, and its gRNA. (A) Schematic diagram of the AYLCB vector constructs expressing CasΦ1, CasΦ2, RiceMINI, and its gRNA under the control of the WDIV-CF promoter. (B) Indel counts and frequencies detected in distal (opposite side of inoculation) and systemically infected leaves of plants inoculated with vectors expressing CasΦ1, CasΦ2, RiceMINI, and its gRNA. (C–E) Representative deletion mutations observed in distal leaves. The gRNA target sequence is highlighted in turquoise, and the PAM sequence is indicated in red. Comprehensive lists of deletions, insertions, and substitutions are provided in Tables S6, S7, and S8.

## Discussion

Most *Mastrevirus* species infect monocots, but some also infect dicots (Liu et al., 1999; Boulton, 2002; Halley-Stott et al., 2007; Nahid et al., 2008). In our previous work, we showed that WDIV, a monocot-infecting *Mastrevirus*, can infect dicots and function as a vector for gene silencing and editing (Kumar et al., 2024). The association of a betasatellite further enhances WDIV’s potential as a vector.

Geminivirus-derived replicons offer efficient delivery of genome editing reagents due to their large cargo capacity and high levels of gene expression (Baltes et al., 2014; Gil-Humanes et al., 2017; Gong et al., 2024). However, their limited systemic movement restricts editing to the inoculation site. To address this, we explored whether a WDIV-based vector—augmented by its associated betasatellite AYLCB—could enable systemic delivery of editing reagents.

We pursued two major goals: (1) develop an AYLCB-based vector by removing the C1 gene; and (2) reduce cargo size by employing viral internal promoters to drive gene expression. The C1 promoter of betasatellite, previously shown to drive constitutive expression of marker genes such as GUS and GFP (Zhang et al., 2012; Shukla et al., 2013), was tested alongside the virion and complementary strand promoters of WDIV using GUS transgenics and AmCyan transient expression. In both assays, the WDIV complementary strand promoter showed the highest activity, followed by CaMV35S, WDIV-VF, and AYLCB-C1—consistent with earlier findings (Xie et al., 2003; Khan et al., 2015; Alok et al., 2019; Zhang et al., 2012; Shukla et al., 2013).

Using these results, we designed CRISPR–Cas cassettes under the control of AYLCB-C1 and WDIV-CF promoters. We then asked whether viral promoters could drive functional gRNA expression. Spacer:gRNA:spacer constructs under the control of viral promoters were delivered using WDIV and AYLCB vectors. AtU6:gRNA constructs were also delivered using WDIV and AYLCB vectors. Indel frequencies, as measured by Cas-Analyzer and CRISPResso2, were comparable across promoters. Notably, the AYLCB-delivered constructs produced the highest indel frequencies, particularly in upper (systemic) leaves. This may be due to the smaller size (∼250 bp) of the spacer:gRNA:spacer cassette compared to the AtU6:gRNA construct (∼550 bp).

Encouraged by these results, we tested co-delivery of Cas9 and its gRNA using a single construct driven by the AYLCB-C1 promoter and flanked by tRNA spacers. Despite Cas9’s large size, we observed indels of up to 6.3% in the inoculated leaf and 1.3–1.5% on the opposite side of the leaf, confirming successful expression and editing.

To overcome the large size limits imposed by the Cas9, we evaluated four miniature Cas variants—CasΦ1, CasΦ2, CasMINI, and RiceMINI—delivered with gRNA using the same strategy. Both Cas-Analyzer and CRISPResso2 detected editing primarily in the inoculated leaf and to a lesser extent in systemic tissue. CasΦ2 showed the highest editing efficiency, followed by CasΦ1 and RiceMINI. CasMINI failed to produce consistent editing, which may benefit from codon optimization.

We then tested whether stronger expression could enhance editing by inserting the WDIV-CF promoter upstream of the C1 promoter in constructs for CasΦ1, CasΦ2, and RiceMINI (excluding CasMINI due to poor performance). Insertion of the WDIV-CF promoter significantly improved editing efficiency. CRISPResso2 consistently predicted higher indel frequencies than Cas-Analyzer, likely due to differences in analytical parameters. For example, sample CP21 yielded 1.20% indels via CRISPResso2 vs. 0.8% via Cas-Analyzer. These differences were consistent across other miniature Cas constructs (Table 1, Table S1). Moreover, the enhanced editing efficiency in the distal tissue suggests that a germline edit is possible if we optimize delivery and editing efficiency.

The PCR-based amplification of the inserts on the viral vectors (such as AmCyan, gRNA, and Cas molecules) showed a lower abundance or absence of delivered DNA in the systemically infected leaves, which reflects vector instability due to the large cargo. Most significantly, we demonstrated tissue culture-free genome editing using a novel viral promoter–tRNA spacer system. AYLCB enabled systemic delivery and expression of both Cas and gRNA. To our knowledge, this is the first report of a geminivirus-based system achieving editing in systemically infected plant tissues. Given the wide host range of WDIV and AYLCB, this platform could facilitate genome editing across diverse plant species without tissue culture. Future work would focus on enhancing editing efficiency, performing templated editing, and germline editing via floral inoculation using this system.

## Materials and Methods

### Vector Development

To develop the AYLCB-based vector, we used a wheat-infecting isolate of AYLCB (clone 72(β1); GenBank accession MN240347) from our previous study (Kumar et al., 2020b) as a template for PCR amplification. The vector was engineered by removing the C1 gene and introducing a multiple cloning site (MCS) downstream of the C1 promoter (Figure S5). Primers P1 and P2 amplified the region containing the satellite conserved region (SCR), A-rich region, and C1 promoter, while primers P3 and P4 amplified part of the SCR and the remaining AYLCB backbone. The PCR products from both sets were digested with *EcoRI* and ligated. This ligation product was used as a template for two additional PCRs using primers P1/P4 and P5/P6, respectively. Both products were cloned into the pJET1.2 vector (Thermo Fisher Scientific, USA), and one positive clone of each was selected.

The clone containing the P1/P4 product was digested with *XbaI* and *AscI*, while the P5/P6 product was digested with *AscI* and *PstI*. In parallel, the pCAMBIA1300 vector was digested with *XbaI* and *PstI*. The two inserts and linearized pCAMBIA1300 were ligated (Figure S5). Primer BetaMCSF/R (Table S10) was used to confirm the presence of the insert in the final construct.

To facilitate replication and systemic movement of the AYLCB-based vector, we co-inoculated plants with an infectious clone of WDIV from our previous study (Kumar et al., 2024). A WDIV-based vector developed previously (Kumar et al., 2024) was also used for comparison of gRNA delivery efficiency. All PCR reactions were performed using Phire Hot Start II DNA Polymerase (Thermo Fisher Scientific, USA).

### Promoter Activity Analysis

Cis-regulatory elements in the AYLCB C1 promoter and the bidirectional promoter of WDIV were analyzed using the PlantCARE database (Lescot et al., 2002). Promoter sequences were further compared to other betasatellites and viral genomes (Zhang et al., 2012; Shukla et al., 2013; Xie et al., 2003; Khan et al., 2015; Alok et al., 2019). To assess promoter activity, the AYLCB C1 promoter and WDIV bidirectional promoters were cloned into pCAMBIA1305, replacing the CaMV 35S promoter, resulting in four constructs: AYLCB-C1, WDIV-VF, WDIV-CF, and CaMV35S (Figure 1A). All constructs were transformed into *Agrobacterium tumefaciens*.

One positive clone from each construct was used for the transformation of *Nicotiana tabacum*, following the protocol described by Jogam et al. (2023). Leaf explants were surface sterilized using bleach and 70% ethanol, washed three times, and cultured on regeneration medium supplemented with 1.0 mg/L BAP, 0.1 mg/L NAA, and 10 mg/L hygromycin. Regenerated shoots were transferred to MS medium containing 1.0 mg/L IAA. All media were solidified with 0.8% agar and adjusted to pH 5.7–5.8 before autoclaving. Plants were grown at 26 ± 2°C with a 16/8 h photoperiod under 50 µmol m ² s ¹ white fluorescent light and 50–60% relative humidity.

At the 4–5 leaf stage, transgenic and wild-type plantlets were subjected to histochemical GUS assays. The GUS solution was prepared from Sigma-Aldrich’s GUS assay kit according to the manufacturer’s instructions. Whole plantlets were incubated overnight in GUS solution at 37°C, followed by chlorophyll removal using 70% ethanol for visualization of GUS expression.

### Development of *AmCyan* Expression Constructs

To evaluate the delivery and expression capabilities of the AYLCB-based vector, we developed four constructs containing the *AmCyan* gene (Wenck et al., 2003) as a visual marker. The first construct was assembled by cloning *AmCyan* and the *Nos* terminator (AN) under control of the AYLCB C1 promoter (Figure 2A). *AmCyan* was amplified using AmCyanKpnFor/AmCyanEcoRev primers, and *Nos* was amplified using NosEcorFor/NosAscRev primers (Table S10). The resulting PCR products were digested with *EcoRI* and ligated. This ligation product served as a template for PCR using AmCyanKpnFor and NosAscRev primers. The final amplicon was digested with *KpnI* and *AscI*, and ligated into the AYLCB vector.

In the second construct series, *AmCyan* was amplified with AmCyanBamFor/AmCyanEcoRev primers, digested with *EcoRI*, and ligated to *EcoRI*-digested *Nos*. The ligation product was used as template for PCR with AmCyanBamFor and NosAscRev primers. The resulting AN fragment was digested with *BamHI* for downstream cloning.

To test different promoters, we amplified the complementary (CF) and virion (VF) strand promoters from WDIV using WDCFBamF/WDCFKpnR and WDVFKpnF/ WDVFBamR primer pairs, respectively. The CaMV35S promoter was amplified using 35sKpnFor/35sBamRev primers. Each promoter fragment was digested with *BamHI* and ligated separately into *BamHI*-digested AN. The resulting ligated products were amplified using primer pairs 35sKpnFor/NosAscRev, WDfullComKpnRev/NosAscRev, and WDfullVirKpnFor/NosAscRev, producing three promoter:gene constructs—35AN, CFAN, and VFAN (Figure 2A). These were then digested with *KpnI* and *AscI* and cloned into the AYLCB vector.

### Expression Analysis Using Real-Time PCR

Leaf samples were collected from GUS-expressing T transgenic tobacco plants and *AmCyan*-construct–inoculated plants. Genomic DNA and total RNA were extracted using DNeasy and RNeasy Plant Mini Kits (QIAGEN, Germany). RNA was reverse transcribed to cDNA and quantified before use.

qRT-PCR was performed using gene-specific primers for *AmCyan*, *GUS*, *hpt*, and AYLCB (Table S10: AmCyanRTFor/Rev, GUSRTFor/Rev, HptRTFor/Rev, AYLCBRTF/R). Reactions were conducted using SYBR Green chemistry on a Roche LightCycler system (Roche Diagnostics International AG, Switzerland). The *Nicotiana tabacum* actin gene served as an internal control for normalization. Relative expression levels were calculated using the 2^−ΔΔCt method. Each qRT-PCR reaction was performed with three biological replicates.

### Development of gRNA Constructs for Cas9-Expressing Tobacco gRNA Construct with AtU6 Promoter

A gRNA targeting the *PDS* gene in *Nicotiana tabacum* (5′-TTGGTAGTAGCGACTCCATG-3′), previously validated (Maher et al., 2020; Kumar et al., 2024), was used to assess genome editing. The AtU6:gRNA cassette from our earlier study (Kumar et al., 2024) was amplified using AtU6AYLCBKpnFor and gRNAterAYLCBAscRev primers (Table S10). The PCR product was cloned into the pJET1.2 vector for sequencing and subsequently subcloned into the AYLCB vector via *KpnI* and *AscI* digestion (Figure S4). A WDIV vector carrying the same AtU6:gRNA cassette (Kumar et al., 2024) was also included for comparison.

### gRNA Construct with tRNA Spacers

To evaluate tRNA-flanked gRNAs, we developed additional constructs in both WDIV and AYLCB vectors. The tRNA-gRNA-tRNA units were amplified using TobPDSspaAscFor/TobPDSspaMluRev (for WDIV) and TobPDSspaKpnFor/TobPDSspaAscRev (for AYLCB) primers (Table S10), using the AtU6:gRNA cassette as the template. PCR products were digested with *AscI/MluI* for cloning in WDIV or *KpnI/AscI* for cloning in AYLCB and ligated into the corresponding vectors (Figure S4). All clones were confirmed by Sanger sequencing.

### Development of Tissue-Culture-Free gRNA and Cas Constructs in AYLCB vector

We generated five CRISPR-Cas constructs in the AYLCB vector under the control of the C1 promoter: (1) Cas9 + gRNA, (2) CasΦ-1 + gRNA, (3) CasΦ-2 + gRNA, (4) CasMINI + gRNA, and (5) Rice-codon-optimized Cas12f (RiceMINI) + gRNA (Figures 4A, 5A, and 6A). Three additional constructs were prepared by inserting the WDIV-CF promoter between the C1 promoter and each Cas variant (Figure 7A). Each cassette was designed with a tRNA spacer on both sides of the gRNA, as shown in Figure S4.

### CRISPR-Cas Constructs with C1 Promoter

The Cas9 construct was generated by amplifying Cas9 (pDIRECT_23A; Cermak et al., 2017) using Cas9KpnFor/Cas9SpeRev, and the gRNA (from Kumar et al., 2024) using Cas9gRSpeFor/Cas9gRAscRev primers (Table S10). Cas9 and gRNA fragments were digested with *KpnI/SpeI* and *AscI/SpeI*, respectively, and cloned into *KpnI/AscI*-digested AYLCB (Figure 4A).

CasΦ-1 and CasΦ-2 constructs were developed similarly. CasΦ-1 and its gRNA were amplified from Pausch et al. (2020) using CasPhi1SPKpnF/CasPhi1SPSpeR and TbPDSCasPhi1SPSpeF/TbPDSCasPhi1SPAscR primers, respectively. CasΦ-2 and its gRNA were amplified with CasPhi2SPAKpnFor/CasPhi2SPAMluRev and TobPDSCasPhi2spaMluFor/TobPDSCasPhi2spaAscRev. Digestion and cloning were performed using *KpnI/SpeI*(CasΦ-1), *AscI/SpeI* (gRNA), and *KpnI/MluI* and *AscI/MluI* (CasΦ-2 and gRNA), respectively (Figure 5A).

CasMINI and RiceMINI constructs were generated as follows. CasMINI and its gRNA were amplified from Xu et al. (2021) using CasMiniSPAKpnFor/CasMiniSPAMluRev and TobPDSMiniSPAMluFor/TobPDSMiniSPAAscRev. For RiceMINI, a codon-optimized gene fragment (Sukegawa et al., 2023) was synthesized (IDT, USA) and amplified using RiceMiniSPcrKpnFor/RiceMiniSPcrSpeRev (Cas) and RiceMINIgRNASpeFor/RiceMINIgRNAAscRev (gRNA). Cas and gRNA fragments were digested (*KpnI/MluI* or *KpnI/SpeI*, *AscI/MluI* or *AscI/SpeI*) and cloned into the AYLCB vector (Figure 6A). Clones were confirmed by Sanger sequencing. Control constructs expressing only the Cas proteins (Cas9, CasΦ-1, CasΦ-2, CasMINI, RiceMINI) were included for mutation analysis.

### CRISPR-Cas Constructs with WDIV-CF Promoter

To assess promoter strength, three additional constructs were prepared by inserting the WDIV-CF promoter upstream of CasΦ-1, CasΦ-2, or RiceMINI (Figure 7A). WDIV-CF was amplified with WDfullComKpnFor/WDfullComKpnRev and digested with *KpnI*. The digested promoter was ligated into *KpnI*-digested Cas clones described above. Five positive clones were sequenced for each construct, and one correctly oriented clone was selected for downstream experiments.

### Agroinoculation

pCAMBIA1300 vector carrying infectious clone of WDIV, WDIV vector with insert and AYLCB vector with insert were transformed into *Agrobacterium tumefaciens* GV3101. One positive clone from each construct was selected and used for inoculation as described previously (Kumar et al. 2014 and 2024). Wild-type *Nicotiana benthamiana* plants at the five or six leaf stage were co-inoculated with the infectious clone of WDIV and AYLCB vector carrying AmCyan expressing cassette (Figure 2A).

Cas9 expressing transgenic tobacco lines were inoculated in four different combinations. In first and second combinations, the AtU6:gRNA cassette (Figure S4A) was delivered using WDIV (Figure S4B), and the AYLCB vector (Figure S4C). In the third and fourth sets, gRNA with tRNA spacer (Figure S4D) was delivered using WDIV (Figure S4E) and AYLCB (Figure S4F) vectors. WDIV was co-inoculated to support replication of the AYLCB based vector.

In the third set of experiment, wild-type tobacco plants were inoculated with AYLCB vector carrying Cas and gRNA constructs (Figure 4A, 5A and 6A, and 7A). In addition, CasMINI, RiceMINI, CasΦ-1, and Cas-2, which only express AYLCB was inoculated as a negative control. WDIV was also co-inoculated to support replication of AYLCB. The plants were observed and tested at 2 to 3 weeks post-inoculation (WPI).

### Detection of WDIV and AYLCB vectors and CRISPR-Cas reagents

Total DNA was isolated from the systemically infected leaves of agroinoculated plants. WDIV was detected using WDCPF/R primers, whereas AYLCB was detected using the BetaC1(-)F/R primers, respectively (Table S10). The delivery of gRNA by WDIV in systemically infected leaves was tested using VIGSMCSF/R primers, whereas the delivery of gRNA and AmCyan by AYLCB was tested using BCMMCSF/R primers (Table S10). In addition, Cas-specific primers, CasMinDet871F/R, RiceMINdetFor/Rev, CasPhi1Det1.2kbF/R, CasPhi2Det1kbF/R and SpCas9For/Rev were also used (Table S10).

### Detection of edits

DNA from the plant samples, which showed the presence of viral vector and editing reagents, was used as a template for amplifying the partial fragment (384 bp) of the NbPDS gene containing the gRNA target region. NbPDS(MutDet)_F1 and NbPDS(MutDet)_R1 primers (Table S10) were used for amplification. All PCRs were performed using Phire Hot Start II DNA polymerase (Thermo Fisher Scientific, USA). The PCR band showing the desired size was cut from the agarose gel, purified, quantified, and outsourced for amplicon sequencing (Azenta Life Sciences, USA). Sequencing data were analyzed using web-based tools, Cas-Analyzer (Park *et al*., 2017) and CRISPResso2 (Clement *et al*., 2019). For detecting Cas9-induced indels, SpCas9 was selected as a nuclease, whereas, for detecting miniature Cas (Casϕ1, 2, CasMINI and RiceMINI) induced indels, FnCpf1 was selected as a nuclease in Cas-Analyzer. The default parameters of Cas-Analyzer, with a minor modification by changing the minimum frequency to one, were used for both Cas9 and miniature Cas molecules.

In CRISPResso2, a default Cas9 setting was used for detecting Cas9-induced indels. Substitutions were ignored to only detect and quantify indels. However, for detecting indels induced by miniature Cas molecules, a custom protocol from a previous study (Gong *et al*., 2024) was used in CRISPResso2. In this protocol, the center of the quantification window was set as 0 nucleotides from the 3’ end of the target sequence, and the quantification window was set to 10 nucleotides with a window size of 30 nucleotides. The reads were mapped to the reference and analyzed for edits with an average read quality of Phred33 scale >20 and single bp quality >10.

## Author Contributions

JK and SFK planned, organized, and supervised the study. SFK obtained the funds. JK, AA, JF, and AS carried out the experiment. JK, AA and FZ analyzed the data and wrote the manuscript draft. JK, FZ, DFV, and SFK revised the manuscript.

## Acknowledgments

Funding for the project was provided by the U.S. Department of Agriculture – Agricultural Research Service (USDA-ARS) National Plant Disease Recovery System (NPDRS). Any opinions, findings, conclusions, or recommendations expressed in this publication are those of the author(s) and do not necessarily reflect the view of the U.S. Department of Agriculture. We thank Roger Casper, USDA-ARS Cereal Disease Lab, for excellent technical assistance. Mention of trade names or commercial products in this publication is solely for the purpose of providing specific information and does not imply recommendation or endorsement by the US Department of Agriculture. USDA is an equal opportunity provider and employer.

## Ethics approval

No ethical approval required as the study did not involve animal/human research.

## References

Acha G, Vergara R, Muñoz M, Mora R, Aguirre C, Muñoz M, Kalazich J et al. (2021) Traceable DNA-Replicon Derived Vector to Speed Up Gene Editing in Potato: Interrupting Genes Related to Undesirable Postharvest Tuber Traits as an Example. Plants, 10, 1882.

Alok A, Tiwari S, Tuli R (2019) A bidirectional promoter from Papaya leaf crumple virus functions in both monocot and dicot plants. Physiol. Mol. Plant Pathol. 108, 0885–5765.

Baltes NJ, Gil-Humanes J, Cermak T, Atkins PA, Voytas DF (2014) DNA replicons for plant genome engineering. Plant Cell, 26, 151–163.

Bigelyte G, Young JK, Karvelis T, Budre K, Zedaveinyte R, Djukanovic V, Van Ginkel E et al. (2021) Miniature type V-F CRISPR-Cas nucleases enable targeted DNA modification in cells. Nat. Commun. 12, 6191.

Boulton MI (2002) Functions and interactions of mastrevirus gene products. Physiol. Mol. Plant Pathol. 60, 243–255.

Catoni M, Noris E, Vaira AM, Jonesman T, Matić S, Soleimani R, Behjatnia SAA et al. (2018) Virus-mediated export of chromosomal DNA in plants. Nat. Commun. 9, 5308.

Cermak T, Curtin SJ, Gil-Humanes J, Cegan R, Kono TJY, Konecna E, Belanto JJ et al. (2017) A multi-purpose toolkit to enable advanced genome engineering in plants. Plant Cell, 29, 1196–1217.

Clement K, Rees H, Canver MC, Gehrke JM, Farouni R, Hsu JY, Cole MA et al. (2019) CRISPResso2 provides accurate and rapid genome editing sequence analysis. Nat. Biotechnol. 37, 224–226.

Gil-Humanes J, Wang Y, Liang Z, Shan Q, Ozuna CV, Sánchez-León S, Baltes NJ et al. (2017) High-efficiency gene targeting in hexaploid wheat using DNA replicons and CRISPR/Cas9. Plant J. 89, 1251–1262.

Gong Z, Previtera DA, Wang Y, Botella JR (2024) Geminiviral-induced genome editing using miniature CRISPR/Cas12j (CasΦ) and Cas12f variants in plants. Plant Cell Rep. 43, 71.

Halley-Stott RP, Tanzer F, Martin DP, Rybicki EP (2007) The complete nucleotide sequence of a mild strain of *Bean yellow dwarf virus*. Arch. Virol. 152, 1237–1240.

Jogam P, Sandhya D, Alok A, Peddaboina V, Singh SP, Abbagani S, Zhang B et al. (2023) Editing of TOM1 gene in tobacco using CRISPR/Cas9 confers resistance to *Tobacco mosaic virus*. Mol Biol Rep. 50, 5165–5176.

Khan ZA, Abdin MZ, Khan JA (2015) Functional characterization of a strong bi-directional constitutive plant promoter isolated from *cotton leaf curl Burewala virus*. PLoS One, 10, e0121656.

Kim DY, Lee JM, Moon SB, Chin HJ, Park S, Lim Y, Kim D et al. (2022) Efficient CRISPR editing with a hypercompact Cas12f1 and engineered guide RNAs delivered by adeno-associated virus. Nat. Biotechnol. 40, 94–102.

Kumar J, Singh SP, Kumar J, Tuli R (2012) A novel mastrevirus infecting wheat in India. Arch. Virol. 157, 2031–2034.

Kumar J, Kumar J, Singh SP, Tuli R (2014) Association of satellites with a mastrevirus in natural infection: complexity of *Wheat dwarf India virus* disease. J. Virol. 88, 7093–7104.

Kumar J, Kumar S, Kianian SF (2020a) The wheat dwarf India virus-betasatellite complex has a wider host range than previously reported. Plant Health Prog. 21, 119–122.

Kumar J, Ahmad J, Imtiaz M, Kianian SF (2020b) *Wheat dwarf India Virus* and associated betasatellite infecting wheat in Pakistan. Austral. Plant Dis. Notes 15, 16.

Kumar J, Alok A, Steffenson BJ, Kianian SF (2024) A geminivirus crosses the monocot-dicot boundary and acts as a viral vector for gene silencing and genome editing. J. Adv. Res. 61, 35–45.

Lescot M, Déhais P, Moreau Y, De Moor B, Rouzé P, Rombauts S (2002) PlantCARE: a database of plant cis-acting regulatory elements and a portal to tools for in silico analysis of promoter sequences. Nucleic Acids Res. 30, 325–327.

Liu L, Saunders K, Thomas CL, Davies JW, Stanley J (1999) *Bean yellow dwarf virus* RepA, but not Rep, binds to maize retinoblastoma protein, and the virus tolerates mutations in the consensus binding motif. Virology, 256, 270–279.

Maher MF, Nasti RA, Vollbrecht M, Starker CG, Clark MD, Voytas DF (2020) Plant gene editing through de novo induction of meristems. Nat. Biotechnol. 38, 84–9.

Mikhaylova E (2025) Virus-Induced Genome Editing (VIGE): One Step Away from an Agricultural Revolution. Int. J. Mol. Sci. 26, 4599.

Nahid N, Amin I, Mansoor S, Rybicki EP, van der Walt E, Briddon RW (2008) Two dicot-infecting mastreviruses (family *Geminiviridae*) occur in Pakistan. Arch. Virol. 153, 1441–1451.

Nasti RA, Voytas DF (2021) Attaining the promise of plant gene editing at scale. Proc Natl Acad Sci USA. 118, e2004846117.

Park J, Lim K, Kim JS, Bae S (2017) Cas-analyzer: an online tool for assessing genome editing results using NGS data. Bioinformatics, 33, 286–288.

Pausch P, Al-Shayeb B, Bisom-Rapp E, Tsuchida CA, Li Z, Cress BF, Knott GJ et al. (2020) CRISPR-CasΦ from huge phages is a hypercompact genome editor. Science, 369, 333–337.

Pausch P, Soczek KM, Herbst DA, Tsuchida CA, Al-Shayeb B, Banfield JF, Nogales E et al. (2021) DNA interference states of the hypercompact CRISPR–CasΦ effector. Nat Struct Mol Biol 28, 652–661.

Shen Y, Ye T, Li Z, Kimutai TH, Song H, Dong X, Wan J (2024) Exploiting viral vectors to deliver genome editing reagents in plants. aBIOTECH. 5, 247–261.

Shukla R, Dalal S, Malathi VG (2013) Isolation and identification of promoter elements of Cotton leaf curl Multan betasatellite associated with Tomato leaf curl viruses. Arch. Phytopathol. Pflanzenschutz, 46, 1015–1029.

Steinberger AR, Voytas DF (2025) Virus-induced gene editing free from tissue culture. Nat. Plants 11, 1241–1251.

Sukegawa S, Nureki O, Toki S, Saika H (2023) Genome editing in rice mediated by miniature size Cas nuclease SpCas12f. Frontiers Genome Edit 5, 1138843.

Takeda SN, Nakagawa R, Okazaki S, Hirano H, Kobayashi K, Kusakizako T, Nishizawa T et al. (2021) Structure of the miniature type V-F CRISPR-Cas effector enzyme. Mol Cell. 81, 558–570.e3.

Wenck A, Pugieux C, Turner M, Dunn M, Stacy C, Tiozzo A, Dunder E et al. (2003) Reef-coral proteins as visual, non-destructive reporters for plant transformation. Plant Cell Rep. 22, 244–51.

Wu, Z., Zhang, Y., Yu, H., Pan, D., Wang, Y., Wang, Y., Li, F., et al. (2021) Programmed genome editing by a miniature CRISPR-Cas12f nuclease. Nat Chem Biol 17, 1132–1138.

Xiao R, Li Z, Wang S, Han R, Chang L (2021) Structural basis for substrate recognition and cleavage by the dimerization-dependent CRISPR-Cas12f nuclease. Nucleic Acids Res. 49, 4120–4128.

Xie Y, Liu Y, Meng M, Chen L, Zhu Z (2003) Isolation and identification of a super strong plant promoter from *cotton leaf curl Multan virus*. Plant Mol Biol. 53, 1–14.

Xu X, Chemparathy A, Zeng L, Kempton HR, Shang S, Nakamura M, Qi LS (2021) Engineered miniature CRISPR-Cas system for mammalian genome regulation and editing. Mol Cell 81, 4333–4345.

Zhang J, Zhang X, Wang Y, Hou H, Qian Y (2012) Characterization of sequence elements from Malvastrum yellow vein betasatellite regulating promoter activity and DNA replication. Virol J. 9, 234.

Zhang C, Liu S, Li X, Zhang R, Li J (2022) Virus-Induced Gene Editing and Its Applications in Plants. Int J Mol Sci. 23, 10202.

