## Supplementary Figures for "A Tissue Culture–Free Genome Editing Strategy in Plants Using Broad-Host-Range Viral Vectors Derived from Geminiviruses"

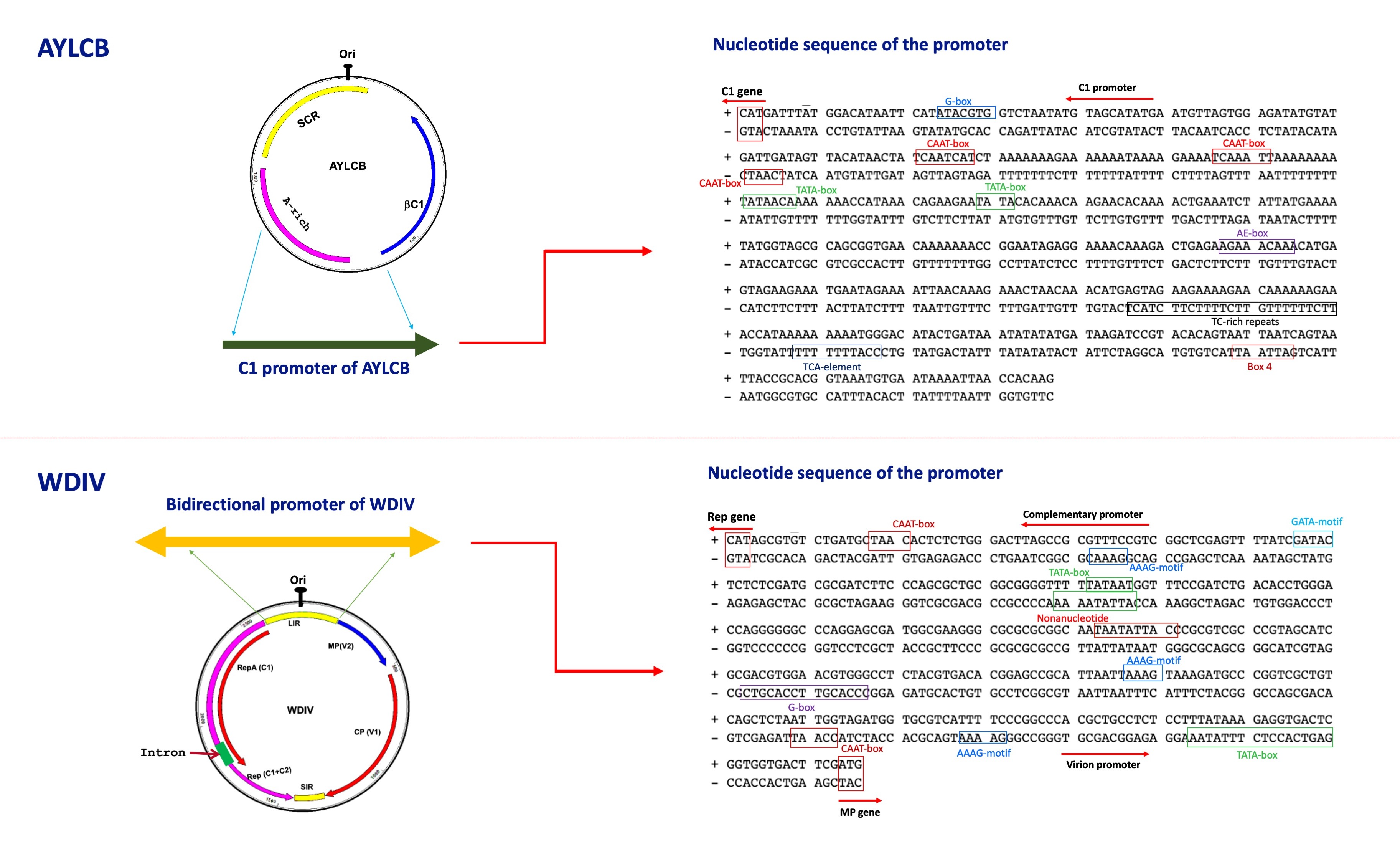

**Figure S1.** Genome organization of Ageratum yellow leaf curl betasatellite (AYLCB) and *Wheat dwarf India virus* (WDIV) representing location of C1 promoter on AYLCB and bidirectional promoter on WDIV are on the left side. The nucleotide sequence of the C1 promoter and bidirectional promoters showing putative cis-acting elements, TATA box, CAAT-box, G-box, GATA-motif, AAAG motif, Box 4, AE- Box, are on the right side. The putative translation start sites ATG and CAT are shown by red arrow and red box at the end of the promoter sequences.

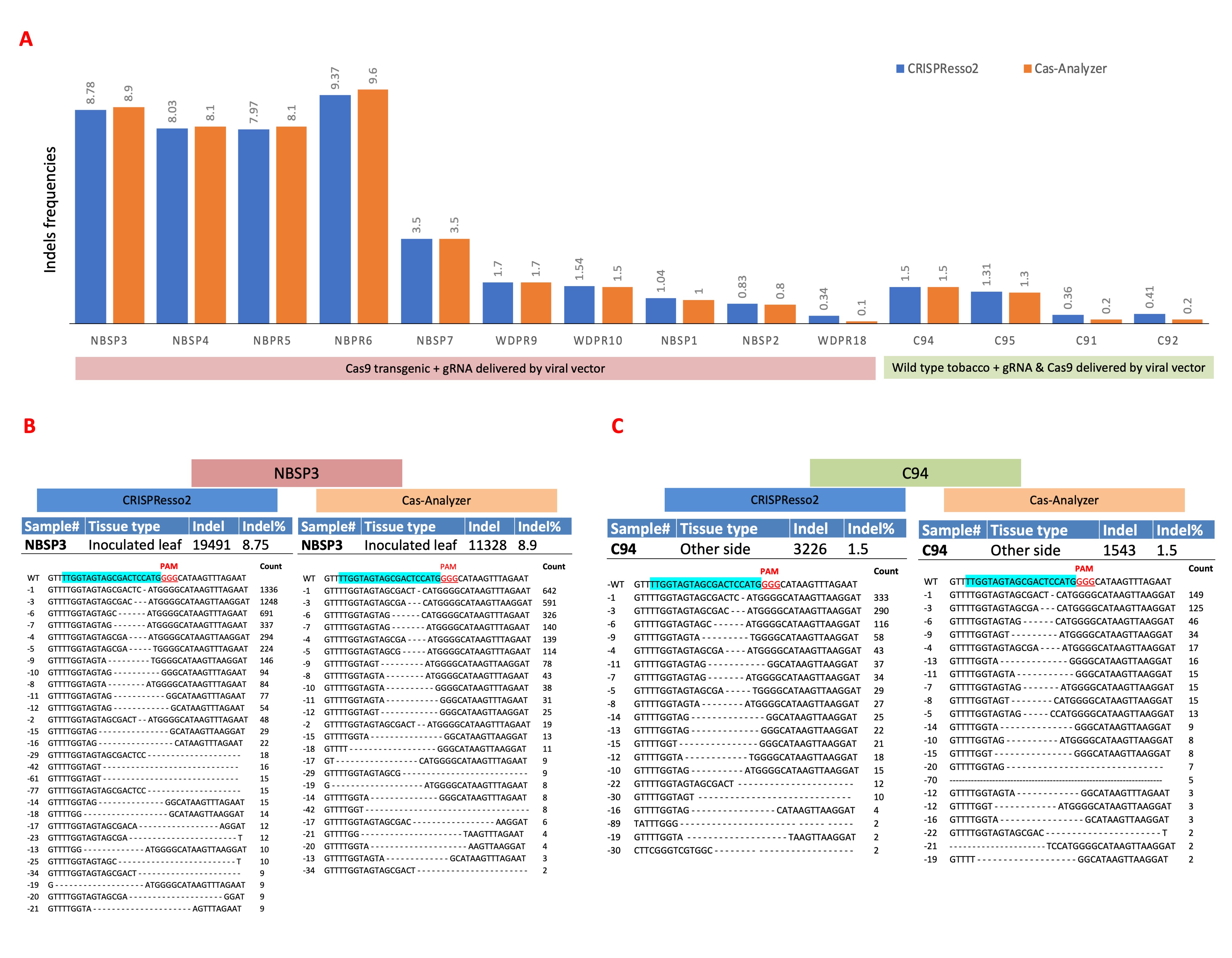

**Figure S2.** Comparison of edit analysis by CRISPResso2 and Cas-Analyzer. (A) Indel frequencies predicted by CRISPResso2 and Cas-Analyzer are compared side by side for each samples having Cas9 induced indels. In spite of having difference in number of indels, both the tools predicted same indel frequencies. (B and C) Representation of a few deletion mutations from the inoculated leaf and systemically infected leaf. The gRNA target sequence is in turquoise color and the PAM region is shown in red color. A more detailed deletion, insertion, and substitution can be found in Table S6, S7 and S8. Both, Cas-Analyzer and CRISPResso2 detected similar deletions but CRISPResso2 predicted a higher count for some of the most prevalent deletions.

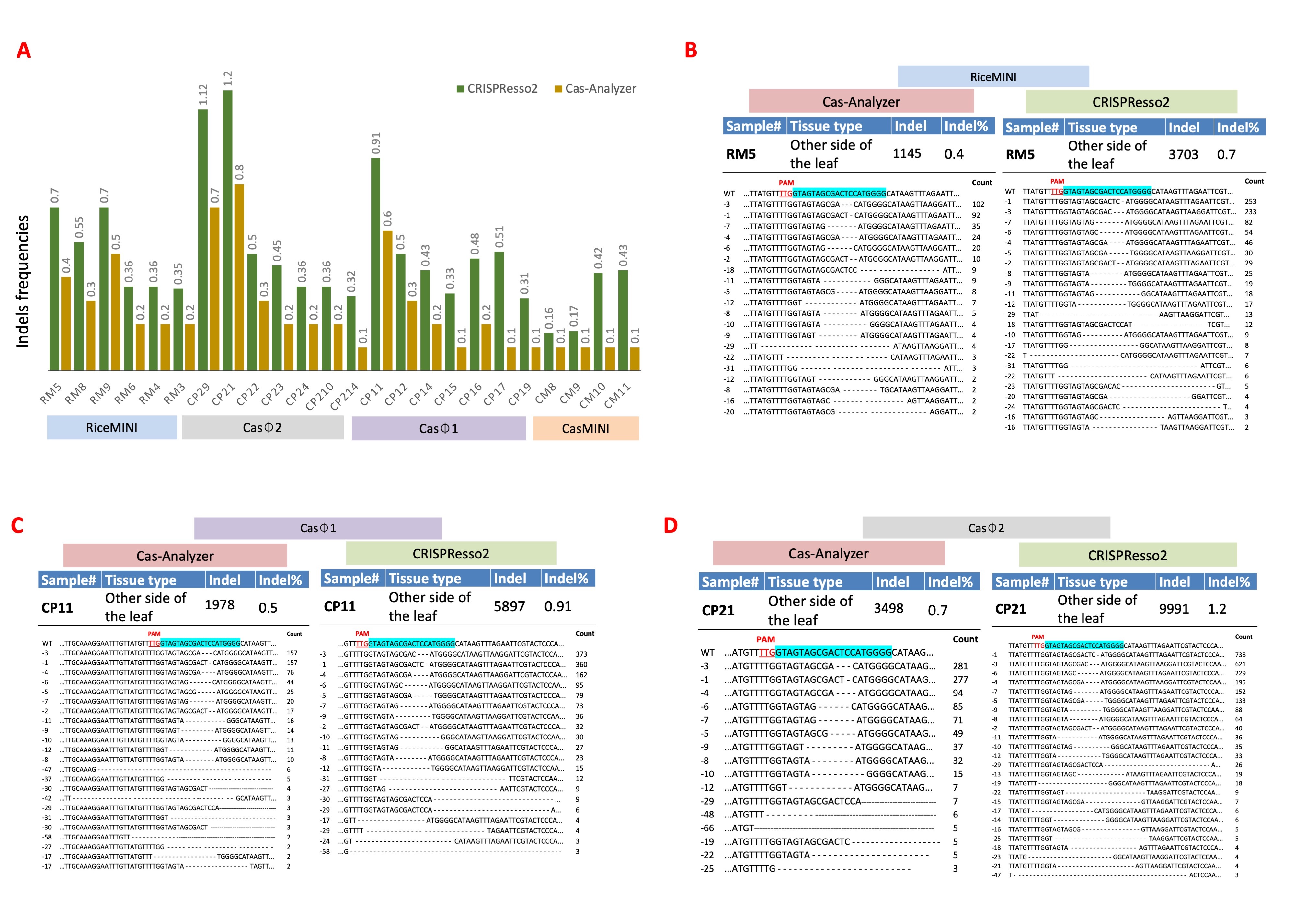

**Figure S3.** Comparison of edit analysis by CRISPResso2 and Cas-Analyzer. (A) Indel frequencies predicted by CRISPResso2 and Cas-Analyzer are compared side by side for each samples having miniature Cas (RiceMINI Casϕ2, Casϕ1 and CasMINI) induced indels. CRISPResso2 predicted a higher indel frequency as compared to Cas-Analyzer. (B-D) Representation of a few deletion mutations from the other side of the leaf. The gRNA target sequence is in turquoise color and the PAM region is shown in red color. Both, Cas-Analyzer and CRISPResso2 detected similar deletions but CRISPResso2 predicted a higher count for some of the most prevalent deletions.

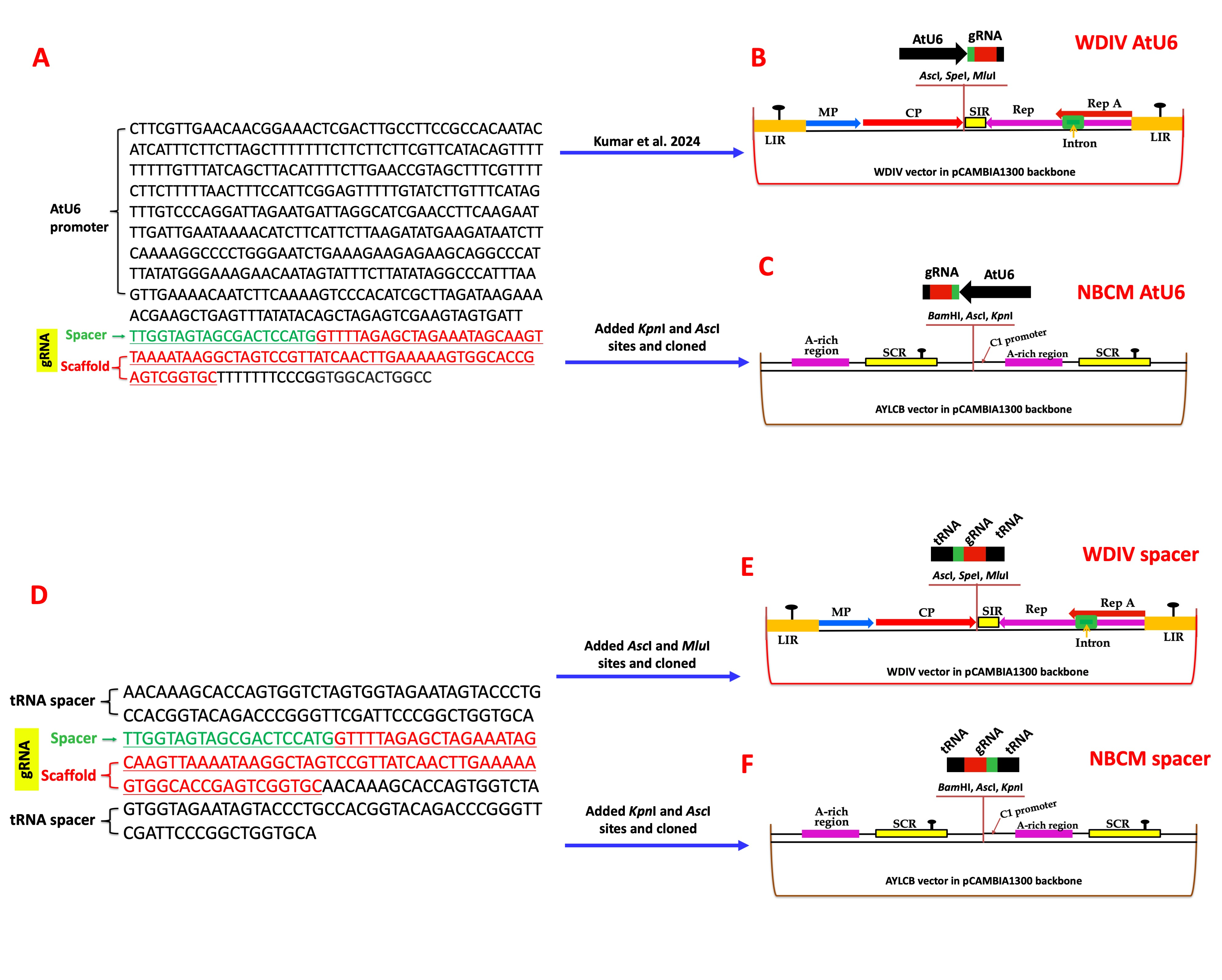

**Figure S4. Schematic representation of Cas9 gRNA constructs.** Nucleotide sequence of AtU6:gRNA (A) and its cloning in WDIV based vector (B) and AYLCB based vector (C). Nucleotide sequence of spacer:gRNA:spacer (D) and its cloning in WDIV based vector (E) and AYLCB based vector (F).

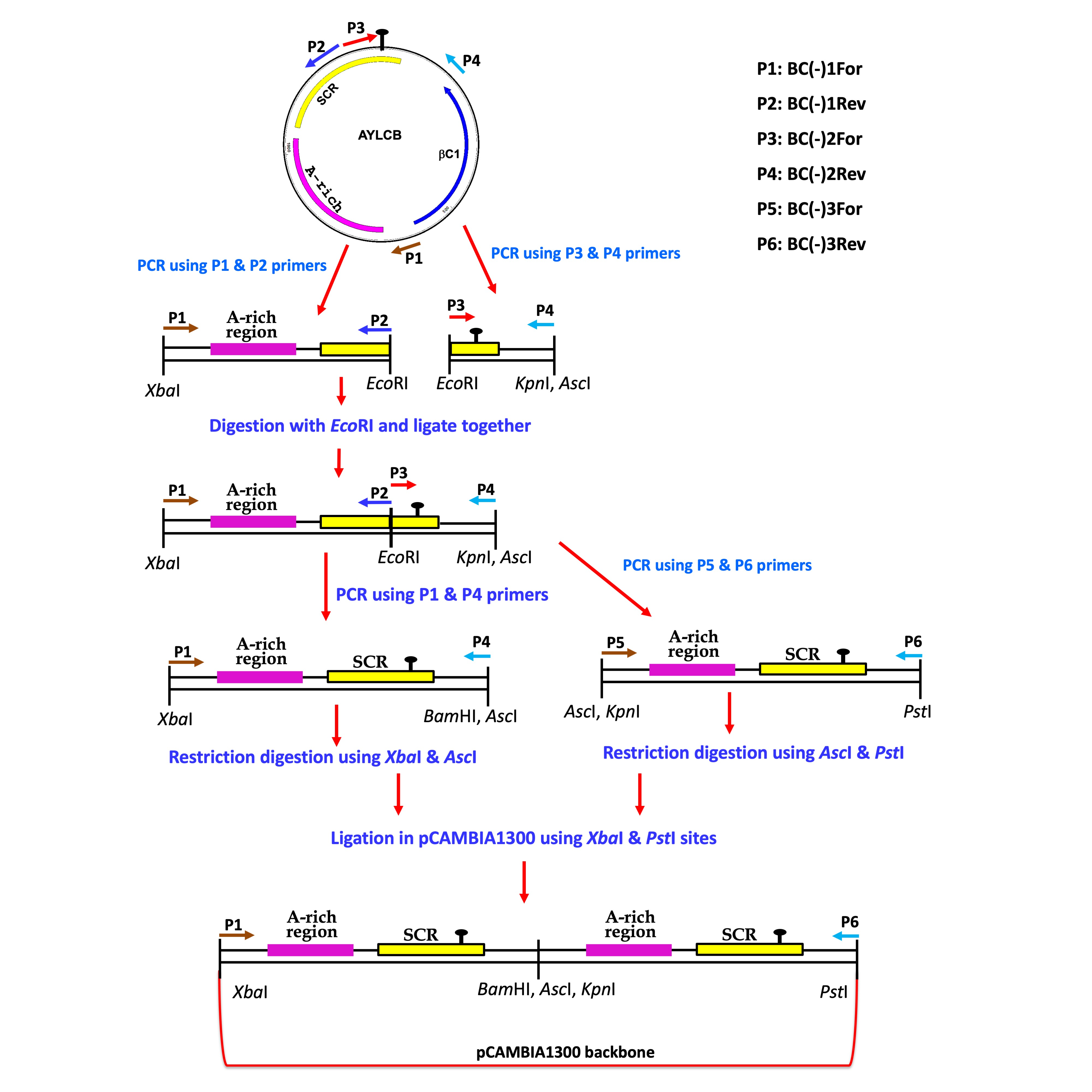

**Figure S5.** Schematic representation of the development of AYLCB-based vector for tissue culture free genome editing. Protocol was followed as mentioned in the flowchart above. Primer details are given in Table S10.
