## Supplementary Table 1 for "A Tissue Culture–Free Genome Editing Strategy in Plants Using Broad-Host-Range Viral Vectors Derived from Geminiviruses"

**Table S1**: Summary of modifications (Indels) in different samples predicted by CRISPResso2

| **Experiment** | **Construct** | **Sample#** | **Tissue type** | **Total reads** | **Reads aligned** | **Indel** | **Indel%** |
| --- | --- | --- | --- | --- | --- | --- | --- |
| Delivery of gRNA in Cas9 transgenic tobacco | NBCM spacer | NBSP3 | Inoculated leaf | 251075 | 221942 | 19491 | 8.78 |
|  |  | NBSP4 |  | 122327 | 105870 | 8500 | 8.03 |
|  | NBCM AtU6 | NBPR5 |  | 122440 | 106921 | 8521 | 7.97 |
|  |  | NBPR6 |  | 129325 | 105206 | 9856 | 9.37 |
|  | NBCM Spacer | NBSP7 | Systemically infected leaf | 851581 | 734222 | 25717 | 3.50 |
|  | WDIV AtU6 | WDPR9 |  | 118435 | 96476 | 1640 | 1.70 |
|  |  | WDPR10 |  | 128967 | 108598 | 1673 | 1.54 |
|  | NBCM spacer | NBSP1 | Top leaf | 112539 | 93449 | 969 | 1.04 |
|  |  | NBSP2 |  | 167013 | 129212 | 1073 | 0.83 |
|  | WDIV AtU6 | WDPR18 |  | 669457 | 569383 | 1912 | 0.34 |
| Delivery of Cas9 and gRNA in wild type tobacco | C1 promoter | C94 | Other side of the leaf | 223701 | 215151 | 3226 | 1.50 |
|  |  | C95 |  | 144481 | 109266 | 1434 | 1.31 |
|  |  | C91 | Next leaf | 685045 | 560782 | 2029 | 0.36 |
|  |  | C92 |  | 654548 | 502322 | 2060 | 0.41 |
| Delivery of RiceMINI and gRNA in wild type tobacco | Comp promoter | RM5 | Other side of the leaf | 707926 | 529719 | 3703 | 0.70 |
|  | C1 promoter | RM8 |  | 742894 | 655236 | 3607 | 0.55 |
|  |  | RM9 |  | 268053 | 248985 | 1749 | 0.70 |
|  | Comp promoter | RM6 | Systemically infected leaf | 800017 | 619183 | 2232 | 0.36 |
|  |  | RM4 |  | 760636 | 658222 | 2348 | 0.36 |
|  | C1 promoter | RM3 |  | 725877 | 591936 | 2046 | 0.35 |
| Delivery of CasPhi2 and gRNA in wild type tobacco | Comp promoter | CP29 | Other side of the leaf | 726972 | 615982 | 6893 | 1.12 |
|  |  | CP21 |  | 893199 | 835124 | 9991 | 1.20 |
|  | C1 promoter | CP22 |  | 791433 | 746147 | 3723 | 0.50 |
|  |  | CP23 |  | 745014 | 683595 | 3051 | 0.45 |
|  | Comp promoter | CP24 | Systemically infected leaf | 732546 | 665062 | 2382 | 0.36 |
|  |  | CP210 |  | 779461 | 695965 | 2495 | 0.36 |
|  | C1 promoter | CP214 |  | 802706 | 691461 | 2203 | 0.32 |
| Delivery of CasPhi1 and gRNA in wild type tobacco | Comp promoter | CP11 | Other side of the leaf | 705274 | 648304 | 5897 | 0.91 |
|  |  | CP12 |  | 730177 | 671042 | 3348 | 0.50 |
|  | C1 promoter | CP14 |  | 718823 | 657928 | 2829 | 0.43 |
|  |  | CP15 |  | 724550 | 616735 | 2043 | 0.33 |
|  | Comp promoter | CP16 | Systemically infected leaf | 668295 | 560967 | 2665 | 0.48 |
|  |  | CP17 |  | 680511 | 491503 | 2520 | 0.51 |
|  | C1 promoter | CP19 |  | 735160 | 558684 | 1738 | 0.31 |
| Delivery of CasMINI and gRNA in wild type tobacco | C1 promoter | CM8 | Other side of the leaf | 446742 | 412238 | 661 | 0.16 |
|  |  | CM9 |  | 492585 | 456565 | 763 | 0.17 |
|  |  | CM10 | infected | 230443 | 177748 | 751 | 0.42 |
|  |  | CM11 |  | 128935 | 108330 | 471 | 0.43 |

NBCM and WDIV spacers are shown in Figure 4F and 4E, whereas NBCM and WDIV AtU6 are shown in Figure 4C and 4B.

C1 promoter is AYLCB promoter, and Comp promoter is complementary promoter from WDIV.
