## Supplementary Table 10 for "A Tissue Culture–Free Genome Editing Strategy in Plants Using Broad-Host-Range Viral Vectors Derived from Geminiviruses"

**Table S10**. List of primers used in this study.

| **Name of primers** | **Nucleotide sequence 5’ to 3’** |
| --- | --- |
| BC(-)1For | CTCTAGAGATTTATGGACATAATTCATATAC |
| BC(-)1Rev | GGAATTCACCCTCCCAGGGGTACACA |
| BC(-)2For | GGAATTCGAAACCACTACGCTACGCA |
| BC(-)2Rev | AGGCGCGCCGGATCCGTGTATATTTATGATATCAATAAA |
| BC(-)3For | AGGCGCGCCGGTACCGATTTATGGACATAATTCATATAC |
| BC(-)3Rev | GCTGCAGGTGTATATTTATGATATCAATAAA |
| BetaMCSF | TATCATAAATATACACGGATCCGGCG |
| BetaMCSR | GACCACGTATATGAATTATGTCCAT |
| AmCyanBamFor | CGGGATCCATGGCCCTGTCCAACAAGTTCA |
| AmCyanEcoRev | GGAATTCTTATGATCTGAGTCCGGAGAAGGGCACCACGGAGGT |
| AmCyanKpnFor | GGGTACCATGGCCCTGTCCAACAAGTTCA |
| NosEcorFor | GGAATTCCGTTCAAACATTTGGCAATAAAGT |
| NosAscRev | AGGCGCGCCCCCGATCTAGTAACATAGATGA |
| 35sKpnFor | GGGTACCGCGTATTGGCTAGAGCAGCT |
| 35sBamRev | CGGGATCCAGAGATAGATTTGTAGAGAGAGA |
| WDCFBamF | CGGGATCCAGCGTGTCTGATGCTAACAC |
| WDCFKpnR | GGGTACCCGAAGTCACCACCGAGTCAC |
| WDVFKpnF | GGGTACCAGCGTGTCTGATGCTAACAC |
| WDVFBamR | CGGGATCCCGAAGTCACCACCGAGTCAC |
| AtU6AYLCBKpnFor | GGGTACCCTTCGTTGAACAACGGAAACTCGAC |
| gRNAterAYLCBAscRev | AGGCGCGCCGGCCAGTGCCACCGGGAAAAAAA |
| TobPDSspaAscFor | AGGCGCGCCAACAAAGCACCAGTGGTCTAGTGGTAGAATAGTACCCTGCCACGGTACAGACCCGGGTTCGATTCCCGGCTGGTGCATTGGTAGTAGCGACTCCATGGTTTT |
| TobPDSspaMluRev | GACGCGTTGCACCAGCCGGGAATCGAACCCGGGTCTGTACCGTGGCAGGGTACTATTCTACCACTAGACCACTGGTGCTTTGTTGCACCGACTCGGTGCCACTTTTTCA |
| TobPDSspaKpnFor | GGGTACCAACAAAGCACCAGTGGTCTAGTGGTAGAATAGTACCCTGCCACGGTACAGACCCGGGTTCGATTCCCGGCTGGTGCATTGGTAGTAGCGACTCCATGGTTTT |
| TobPDSspaAscRev | AGGCGCGCCTGCACCAGCCGGGAATCGAACCCGGGTCTGTACCGTGGCAGGGTACTATTCTACCACTAGACCACTGGTGCTTTGTTGCACCGACTCGGTGCCACTTTTTCA |
| Cas9KpnFor | GGGTACCAACAAAGCACCAGTGGTCTAGTGGTAGAATAGTACCCTGCCACGGTACAGACCCGGGTTCGATTCCCGGCTGGTGCAATGGATAAGAAGTACTCTATCGGACT |
| Cas9SpeRev | GACTAGTTCAAACCTTCCTCTTCTTCTTAGGAT |
| Cas9gRSpeFor | GACTAGTAACAAAGCACCAGTGGTCTAGTGGTAGAATAGTACCCTGCCACGGTACAGACCCGGGTTCGATTCCCGGCTGGTGCATTGGTAGTAGCGACTCCATGGTTTT |
| Cas9gRAscRev | AGGCGCGCCTGCACCAGCCGGGAATCGAACCCGGGTCTGTACCGTGGCAGGGTACTATTCTACCACTAGACCACTGGTGCTTTGTTGCACCGACTCGGTGCCACTTTTTCA |
| CasMiniSPAKpnFor | GGGTACCAACAAAGCACCAGTGGTCTAGTGGTAGAATAGTACCCTGCCACGGTACAGACCCGGGTTCGATTCCCGGCTGGTGCAATGGGACCCAAGAAAAAACGCAA |
| CasMiniSPAMluRev | GACGCGTTCAGGCCGGTTCCTCTTTGGT |
| TobPDSMiniSPAMluFor | GACGCGTAACAAAGCACCAGTGGTCTAGTGGTAGAATAGTACCCTGCCACGGTACAGACCCGGGTTCGATTCCCGGCTGGTGCAGCTTCACTGATAAAGTGGAGAA |
| TobPDSMiniSPAAscRev | AGGCGCGCCTGCACCAGCCGGGAATCGAACCCGGGTCTGTACCGTGGCAGGGTACTATTCTACCACTAGACCACTGGTGCTTTGTTCCCCATGGAGTCGCTACTAC |
| RiceMiniSPcrKpnFor | GGGTACCAACAAAGCACCAGTGGTCTAGTGGTAGAATAGTACCCTGCCACGGTACAGACCCGGGTTCGATTCCCGGCTGGTGCAATGAAGAGGACAGCCGACGGCT |
| RiceMiniSPcrSpeRev | GACTAGTTCACTTGGCCTTGCGGATCTTCTT |
| RiceMINIgRNASpeFor | GACTAGTAACAAAGCACCAGTGGTCTAGTGGTAGAATAGTACCCTGCCACGGTACAGACCCGGGTTCGATTCCCGGCTGGTGCAATTTACTCTGTTTCGCGCGCC |
| RiceMINIgRNAAscRev | AGGCGCGCCTGCACCAGCCGGGAATCGAACCCGGGTCTGTACCGTGGCAGGGTACTATTCTACCACTAGACCACTGGTGCTTTGTTAATTGCCCTTCGAAGGGACAAAAAAA |
| CasPhi1SPKpnF | GGGTACCAACAAAGCACCAGTGGTCTAGTGGTAGAATAGTACCCTGCCACGGTACAGACCCGGGTTCGATTCCCGGCTGGTGCAATGGCGGACACACCAACACTTTTC |
| CasPhi1SPSpeR | GACTAGTTCACTTATCATCATCATCCTTGTAGTC |
| TbPDSCasPhi1SPSpeF | GACTAGTAACAAAGCACCAGTGGTCTAGTGGTAGAATAGTACCCTGCCACGGTACAGACCCGGGTTCGATTCCCGGCTGGTGCAGAAACACCGGGAGAGATCTCAAACGATTGCTCGATTAGTCGA |
| TbPDSCasPhi1SPAscR | AGGCGCGCCTGCACCAGCCGGGAATCGAACCCGGGTCTGTACCGTGGCAGGGTACTATTCTACCACTAGACCACTGGTGCTTTGTTCCCCATGGAGTCGCTACTACTCGACTAATCGAGCAATCGT |
| CasPhi2SPAKpnFor | GGGTACCAACAAAGCACCAGTGGTCTAGTGGTAGAATAGTACCCTGCCACGGTACAGACCCGGGTTCGATTCCCGGCTGGTGCAATGCCAAAGCCAGCCGTGGA |
| CasPhi2SPAMluRev | GACGCGTCTTATCATCATCATCCTTGTAGTC |
| TobPDSCasPhi2spaMluFor | GACGCGTAACAAAGCACCAGTGGTCTAGTGGTAGAATAGTACCCTGCCACGGTACAGACCCGGGTTCGATTCCCGGCTGGTGCAGTCGGAACGCTCAACGATTGCCCCTCACGAGGGGAC |
| TobPDSCasPhi2spaAscRev | AGGCGCGCCTGCACCAGCCGGGAATCGAACCCGGGTCTGTACCGTGGCAGGGTACTATTCTACCACTAGACCACTGGTGCTTTGTTCCCCATGGAGTCGCTACTACGTCCCCTCGTGAGGGGC |
| WDfullComKpnFor | GGGTACCAGCGTGTCTGATGCTAACAC |
| WDCPF | ATGTCTCAGGTGAAGAAGAGG |
| WDCPR | TTACTGGTTACCGATACTCTTG |
| BetaC1(-)F | GATTTATGGACATAATTCATAT |
| BetaC1(-)R | GTGTATATTTATGATATCAATAAA |
| NbPDS(MutDet)_F1 | CTTATCTTTGGAGCTCGAGGTCTTC |
| NbPDS(MutDet)_R1 | ACCTGCACCAGCAATAACAATCTCC |
| VIGSMCSF | CAAGAGTATCGGTAACCAGTAA |
| VIGSMCSR | GTTTATTACATATATGAGCAACCT |
| SpCas9For | TCGAGAAGATGGATGGAACC |
| SpCas9Rev | CATTCCCTCGGTCACGTACT |
| BCMMCSF | TTGGGCTTTGAATGGAAT |
| BCMMCSR | GCTACATATTAGACCACGT |
| CasMinDet871F | GTGGTCAGTTTCCGGATGCAGT |
| CasMinDet871R | TGACCTGCACGTTTATGACGGT |
| RiceMINdetFor | CATGAGCAAGGACCACATCG |
| RiceMINdetRev | TACGGGGTGCTGATGTTCTT |
| CasPhi1Det1.2kbF | GCGCTGGATGAGACCGGTCA |
| CasPhi1Det1.2kbR | CCCTTCCGTGAAAGAAGCCGAT |
| CasPhi2Det1kbF | TTCGCCGAGTGGCCAATTATGA |
| CasPhi2Det1kbR | GGTTCACATTGAAGTGCCCCGT |
| BetaC1ProHindFor | CAAGCTTCCTTGTGGTTAATTTTATTCACAT |
| BetaC1ProNcoRev | CCCATGGGATTTATGGACATAATTCATATAC |
| WDfullComF | CCCATGGAGCGTGTCTGATGCTAACAC |
| WDfullComR | CAAGCTTCGAAGTCACCACCGAGTCAC |
| WDfullVirF | CAAGCTTAGCGTGTCTGATGCTAACAC |
| WDfullVirR | CCCATGGCGAAGTCACCACCGAGTCAC |
| HptRTFor | GGAATCGGTCAATACACTACATGGC |
| HptRTRev | AGTCCTCGGCCCAAAGCATCAG |
| GUSRTFor | TGTACAGCGAGCGCCACGAAGAG |
| GUSRTRev | CCCGGTTGGGCCATTGAAGTCGG |
| AmCyanRTFor | TACAAGACCAAGAAGCCCGT |
| AmCyanRTRev | GATCTGAGTCCGGAGAAGGG |
| TobActFor | ATGATATGGAGAAGATCTGGCATCA |
| TobActRev | AGCCTGAATAGCAACATACATAGC |
| AYLCBRTF | TCACCGATTTGATCACCGAT |
| AYLCBRTR | TGATGACCGGAAGTGCCTTA |
